# A mesoscale connectome-based model of conscious access in the macaque monkey

**DOI:** 10.1101/2022.02.20.481230

**Authors:** Ulysse Klatzmann, Sean Froudist-Walsh, Daniel P. Bliss, Panagiota Theodoni, Jorge Mejías, Meiqi Niu, Lucija Rapan, Daniel S. Margulies, Nicola Palomero-Gallagher, Claire Sergent, Stanislas Dehaene, Xiao-Jing Wang

**Author notes:** co-first authors (Ulysse Klatzmann, Sean Froudist-Walsh).

## Abstract

A growing body of evidence suggests that conscious perception of a sensory stimulus coincides with all-or-none activity across multiple cortical areas, a phenomenon called ‘ignition’. In contrast, the same stimulus, when undetected, induces only transient activity. In this work, we report a large-scale model of the macaque cortex based on recently quantified structural mesoscopic connectome data. We use this model to simulate a detection task, and demonstrate how a dynamical bifurcation mechanism produces ignition-like events in the model network. The model predicts that feedforward excitatory transmission is primarily mediated by the fast AMPA receptors to ensure rapid signal propagation from sensory to associative areas. In contrast, a greater proportion of the inter-areal feedback projections and local recurrent excitation depend on the slow NMDA receptors, to ensure ignition of distributed frontoparietal activity. Our model predicts, counterintuitively, that fast-responding sensory areas contain a higher ratio of NMDA to AMPA receptors compared to association cortical areas that show slow, sustained activity. We validate this prediction using cortex-wide *in-vitro* receptor autoradiography data. Finally, we show how this model can account for various behavioral and physiological effects linked to consciousness. Together, these findings clarify the neurophysiological mechanisms of conscious access in the primate cortex and support the concept that gradients of receptor densities along the cortical hierarchy contribute to distributed cognitive functions.

## Introduction

Among the huge flow of information received by our sensory organs, only a fraction of it is consciously perceived (James 1890). The network, cellular and synaptic mechanisms of conscious perception are hotly debated, and largely unresolved (Aru et al. 2020; Baars 2005; Dehaene et al. 1998; Lamme and Roelfsema 2000; Lau and Rosenthal 2011; Mashour et al. 2020; Tononi 2004). In many experiments that probe the access of stimuli to consciousness, subjects (human or non-human) are presented with faint stimuli, and asked to report if they detect them. Neural activity in early sensory areas grows approximately linearly with stimulus strength, regardless of whether the stimulus is detected or not (de Lafuente and Romo 2005, 2006; Del Cul et al. 2007; Kouider et al. 2013; Romo and Rossi-Pool 2020; Van Vugt et al. 2018). However, when a stimulus is consciously detected, activity commonly emerges across frontal and parietal cortex in all-or-none fashion, and is sustained for a few hundred milliseconds (de Lafuente and Romo 2005, 2006; Dehaene et al. 2001; Del Cul et al. 2007; King and Dehaene 2014; Rees et al. 2002; Sadaghiani et al. 2009; Sergent et al. 2005; Van Vugt et al. 2018) in stable (Schurger et al. 2015) or reliable dynamic trajectories (Baria et al. 2017; Salti et al. 2015). This widely distributed, sudden, and sustained activity has been termed ’ignition’ (Dehaene et al. 1998, 2003).

Several prominent theories of consciousness propose a central role of recurrent synaptic interactions between excitatory neurons (Dehaene et al. 2003; Lamme and Roelfsema 2000). However, the timescale of excitatory synaptic interactions differs drastically depending on the type of postsynaptic glutamatergic receptors. The most widely expressed of these are *α*-amino-3-hydroxy-5-methyl-4-isoxazolepropionic acid (AMPA) and N-methyl-D-aspartate (NMDA) receptors. Experimental and theoretical work on working memory has emphasised the importance of AMPA receptors to rapid responses in early sensory cortex and NMDA receptors to sustaining activity in prefrontal cortex (Wang et al. 2013; Wang 1999; Yang et al. 2018). However, studies of conscious perception have hypothesized a critical role of NMDA receptors at long-distance feedback connections (which has been partially supported experimentally, Self et al. 2012). This may imply a large proportion of NMDA receptors in major targets of feedback connections, such as in early sensory cortex, seemingly in contrast to work from the working memory literature. It is unclear whether these two positions are compatible, or if the ignition phenomenon is critically dependent on the synapses at which AMPA and NMDA receptors are expressed.

Small-scale simulations of brain areas organised in perfect hierarchy with NMDA-mediated feedback connections have reproduced the late sustained activity (Dehaene et al. 2003; Dehaene and Changeux 2005; Van Vugt et al. 2018). Though critical for building intuitions, by abstracting away from anatomy, small-scale simulations limit the anatomical specificity of predictions and may overlook major problems that the brain must overcome to enable ignition, such as propagation of information through a strongly-recurrent large-scale system (Joglekar et al. 2018). Large-scale modeling studies based on real cortical connectivity data have investigated the dynamical propagation of stimulus information into the fronto-parietal network, but have not captured the late sustained activity that is seen experimentally (Chaudhuri et al. 2015; Deco et al. 2020; Joglekar et al. 2018). A realistic large-scale model of the ignition phenomenon should create testable, anatomically-precise predictions for theories of consciousness, reproduce key physiological findings from detection experiments across sensory and association cortex and behaviour and offer a platform for future simulations of other experimental paradigms.

In this paper, we develop a mesoscale connectome-based dynamical model of the macaque cortex with realistic biophysical constraints and assess its behavior during a stimulus detection task, similar to that used experimentally. Secondly, we examine whether the parameter regime necessary for realistic model behavior is consistent with receptor distributions in the macaque cortex. Our model reproduces multiple aspects of monkey behavior and physiology, including aspects that have evaded previous models such as strong propagation of activity through the connectome to prefrontal cortex, bifurcation dynamics and sustained activity across a distributed subsets of fronto-parietal regions. Furthermore, we demonstrate that sufficiently strong stimulus propagation and ignition requires NMDA/AMPA distributions across cortex that closely match those measured experimentally. Therefore our findings shed light on the synaptic and systems-level mechanisms underlying ignition and reconcile seemingly contradictory anatomical, physiological and modeling results.

## Results

### A large-scale dynamical model of the macaque cortex with NMDA, AMPA and GABA receptors

We built a large-scale model of the macaque cortex containing 40 different interacting cortical areas. Each cortical area contains a local circuit, with two populations of excitatory neurons and one population of inhibitory neurons (Mejias and Wang 2022; Wong and Wang 2006). Excitatory connections are mediated by both NMDA and AMPA receptors, and inhibitory connections are mediated by *GABA*_*A*_ receptors. Cortical areas differ in the strength of excitatory input (due to differences in the expression of dendritic spines Elston 2007), and are connected according to weighted and directed inter-areal connections, measured by retrograde tract-tracing (Markov et al. 2012) with data from 40 interconnected areas (Froudist-Walsh et al. 2021).

### Stimulus detection is accompanied by widespread ignition of activity throughout the fronto-parietal network

We simulated a stimulus detection task by injecting differing, small amounts of external current to primary visual cortex (V1) for 50ms (Fig 1A). In V1, average activity over trials increased approximately linearly with stimulus intensity, before returning to baseline a few milliseconds after the stimulus was removed (Fig 1B). In contrast, on many trials activity in areas throughout the prefrontal and parietal cortices reached a high activity state at around 200ms (Fig 1B,C). This activity remained stable until the end of the trial, or until the vigilance signal was removed. This pattern of prefrontal and early visual activity closely matches the dynamics of neural activity in monkey prefrontal cortex during a similar task (Van Vugt et al. 2018). We therefore interpret trials with late, sustained activity in area 9/46d of the dorsolateral prefrontal cortex as corresponding to detection of the stimulus (Van Vugt et al. 2018) (i.e. hit trials), and trials without such activity as miss trials.

**Figure 1:**
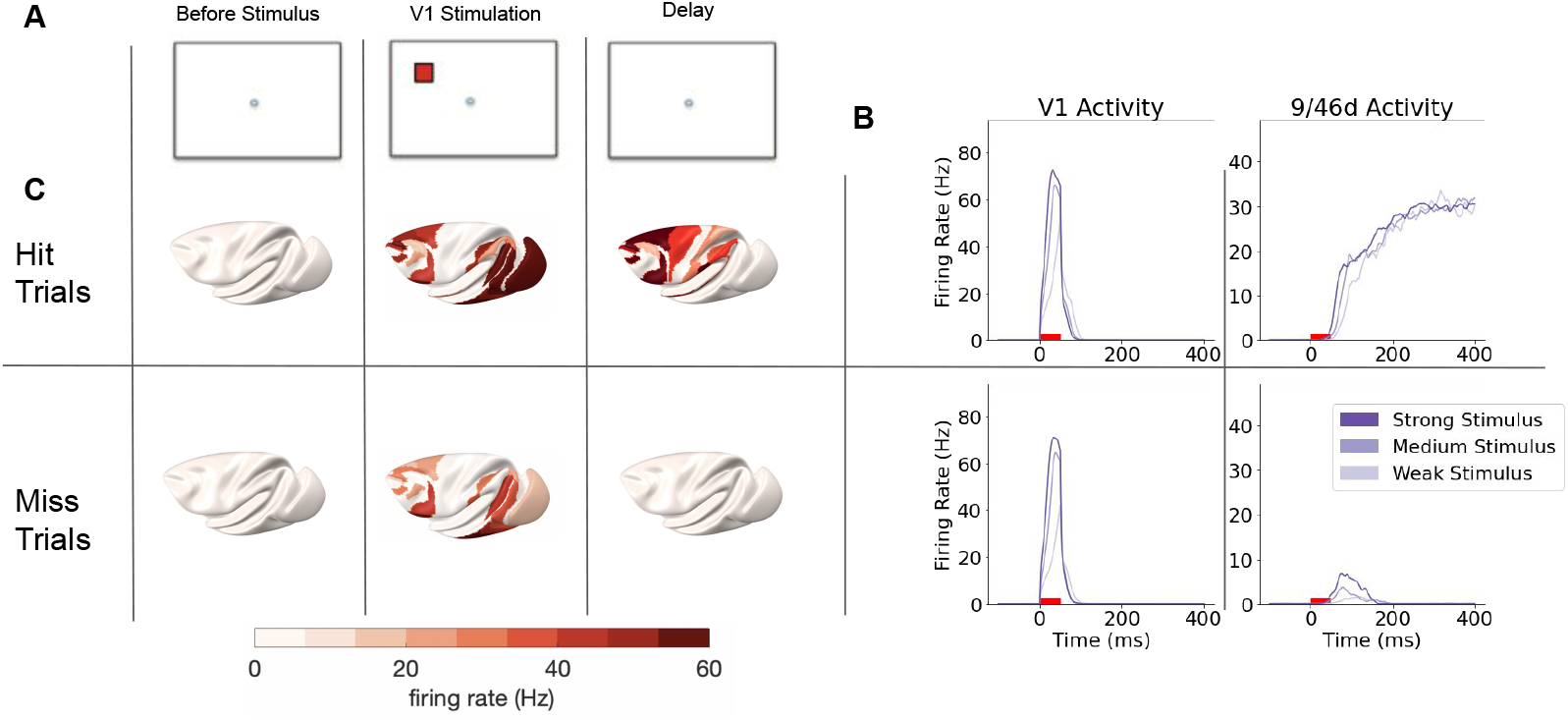
Stimulus detection is accompanied by widespread ignition of activity throughout the fronto-parietal network A) Task structure. A near-threshold stimulus is presented to the excitatory population in area V1 for 50ms. B) Top: Averaged activity over trials in V1 (primary visual cortex) and area 9/46d (prefrontal cortex) during hit trials, for differing levels of stimulus intensity (200pA, 250pA and 300pa respectively for Weak, Medium and Strong stimulus strengths). V1 activity rapidly increases to a peak that differs according to stimulus intensity, before falling back to baseline. 9/46d activity in contrast reaches a high sustained activity state after about 200ms, which does not depend on stimulus intensity. Bottom: Averaged activity over trials in V1 and area 9/46d during miss trials, for differing levels of stimulus intensity. V1 activity is very similar to that on hit trials. 9/46d activity differs drastically, with a smaller peak of activity followed by a return to a low firing baseline state. C) Firing rates across the cortex during example hit (Top) and miss (Bottom) trials. Hit trials are accompanied by sustained activity throughout much of prefrontal and posterior parietal cortex, which is absent on miss trials. Note that in B) and C) stimulus intensity and network parameters are completely matched between hit and miss trials, which differ only in the random noise.

### Activity in the fronto-parietal network, but not sensory areas distinguishes hit from miss trials

Average activity over trials in sensory areas was very similar for hit (stimulus present and detected) and miss (stimulus present but not detected) trials (Fig 1B). Therefore, regardless of whether the stimulus was detected or not, neural activity in sensory areas reliably tracked the objective stimulus strength. In contrast, while hit trials engaged strong, sustained activity throughout frontal and parietal cortices, miss trials led to either a transient increase in activity, which returned to baseline, or no increase in activity (Fig 1B). The sensory and prefrontal findings closely correspond to experimental findings from visual and somatosensory detection experiments (de Lafuente and Romo 2005; Romo and Rossi-Pool 2020; Van Vugt et al. 2018), validating the model. Beyond primary sensory cortex and dlPFC, our model therefore predicts that when a stimulus is detected late stimulus-related activity should be detectable throughout a distributed prefrontal and posterior parietal cortical network (Fig 1C).

### The probability of detecting a stimulus increases nonlinearly with stimulus intensity

Due to the stochastic single-trial behavior, it is possible to analyze how the proportion of hit trials varies with stimulus intensity. Note that late activity always proceeded to either a high or low activity state, as seen in monkey and human experiments (Sergent et al. 2021; Van Vugt et al. 2018). The proportion of hits increased with the stimulus intensity, with a sigmoidal curve accurately fitting the data (Fig. 2), as has frequently been observed in monkeys and humans (de Lafuente and Romo 2006; Del Cul et al. 2007; Van Vugt et al. 2018).

**Figure 2:**
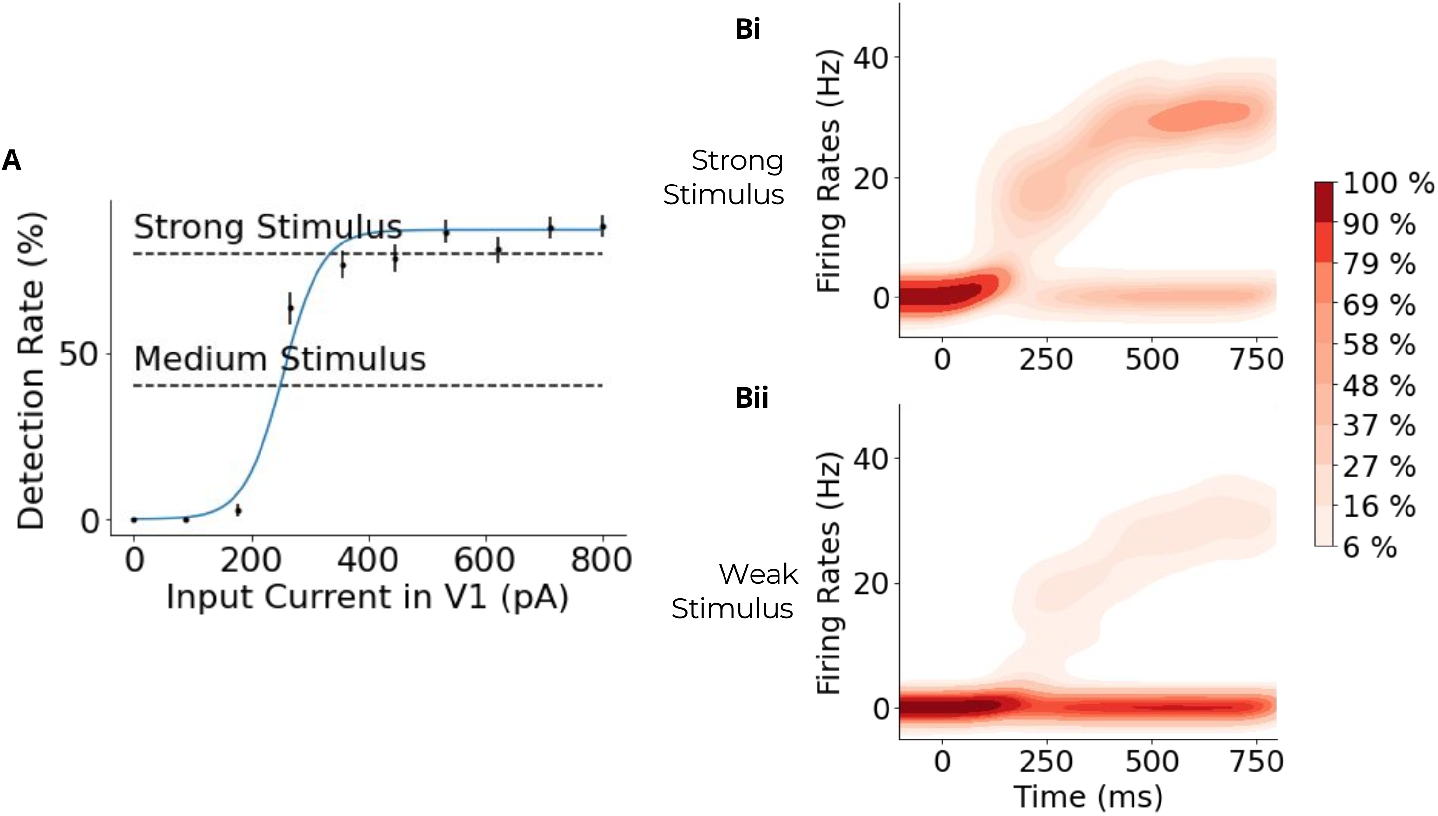
A sigmoidal relationship between stimulus intensity and detection probability. A) The rate at which the large-scale cortical model detects the stimulus (engages sustained activity over 15Hz in area 9/46d) increases non-linearly with the stimulus intensity (input current in V1). B) The distributions of firing rates across trials for area 9/46d in the large-scale model for strong (about 80% detection rate) and weak (about 20% detection rate) stimuli. The outcome of individual trials is stochastic, depending on the noise in the system, but the system always ends in either a high activity state, corresponding to stimulus detection, or a low activity state, corresponding to a miss. A higher percentage of trials with a strong stimulus end in the high activity state compared to trials with a weak stimulus (as seen by the darker red in the high activity branch after 200ms). Stimulus is presented at 0ms for 50ms.

### Transition from early unimodal to late bimodal neural activity in stimulus detection

During stimulus detection tasks, early (*<* 200*ms*) cortical activity increases with increasing stimulus strength, irrespective of whether the stimulus is later detected or missed. When plotted across trials, this early activity creates a unimodal distribution (Sergent et al. 2021), and likely corresponds to pre-conscious processing (Dehaene et al. 2006). After *∼* 250ms, activity either increases to a high activity state, or returns to a low activity state (Sergent et al. 2021), thus creating a bimodal cross-trial distribution, with only the high activity state corresponding to the conscious detection of the stimulus. Thus, Sergent and colleagues suggest that trial activity proceeds from a dynamic sequence of early states to one of two possible late activity states (for prior evidence, see e.g. (Sergent and Dehaene 2003)). This may hint at a bifurcation (i.e. a change in the number or stability of internal states) occurring over time.

Following the methods of Sergent et al., we analyzed model activity at each timepoint across several trials, and examined whether activity across trials was best described by a null distribution (neural activity independent of the stimulus), unimodal distribution or a bimodal distribution. Shortly following the stimulus, activity was best described by a unimodal distribution. After approximately 100ms, the data were best described by a bimodal distribution (Fig 3A), matching experimental observations in humans detecting auditory stimuli (Sergent et al. 2021). In our model, we detect shift to a bimodal cross-trial distribution after about 100ms, with late activity reaching its peak in prefrontal cortex after 200ms (see Fig S1). This broadly matches the timing of stimulus-induced prefrontal activity observed in monkey experiments, which is observable from *∼* 60*ms* following stimulus onset and peaks after *∼* 150 *−* 200*ms* (Bellet et al. 2022; Thorpe and Fabre-Thorpe 2001; Van Vugt et al. 2018). Therefore, our connectome-based dynam-ical model accounts for the temporal progression of activity states observed in the brain during stimulus detection tasks.

**Figure 3:**
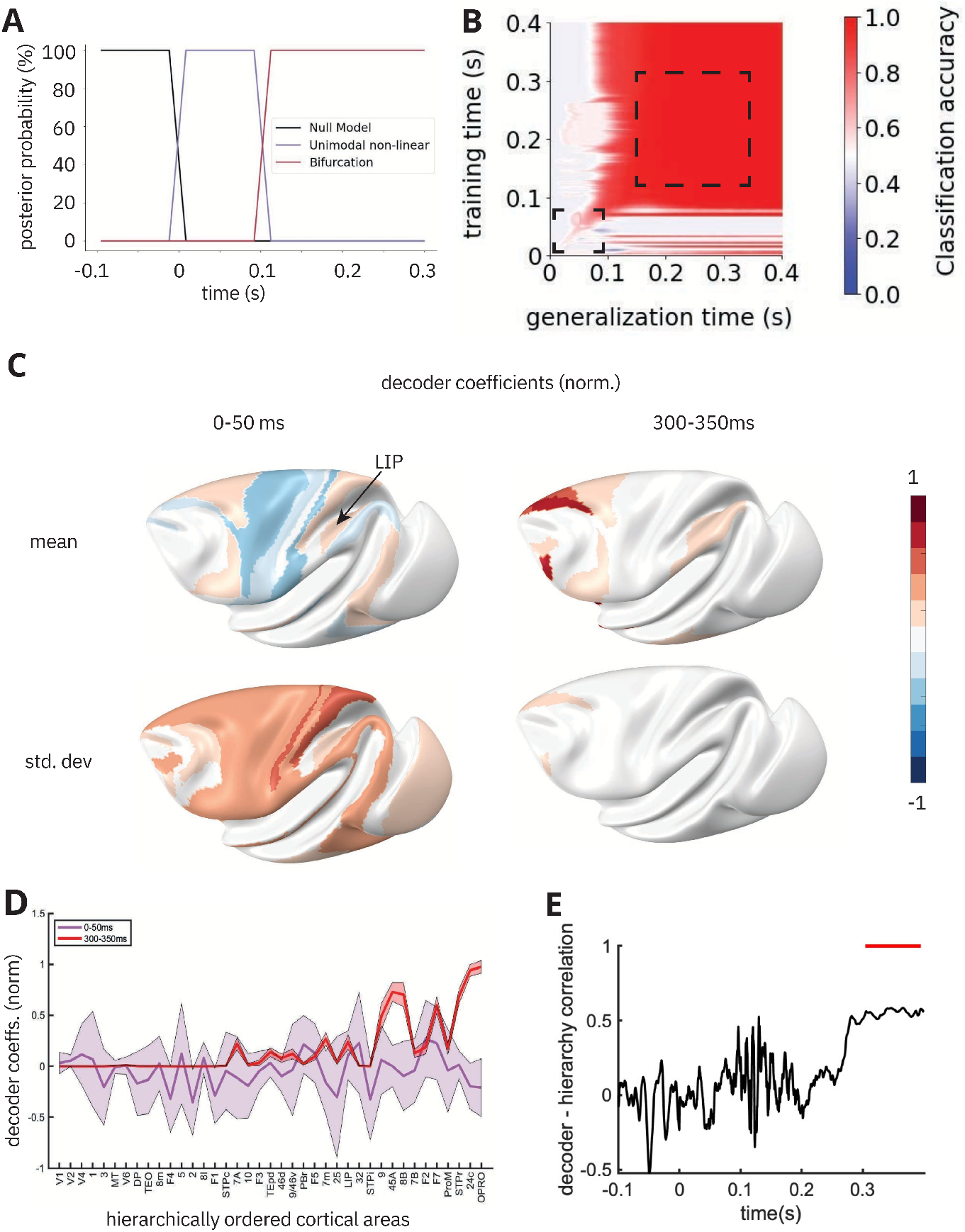
A dynamic-to-sustained progression of activity states associated with ignition. A) Across-trial statistics of neural activity for different stimulus strengths were used to classify model neural activity as belonging to null (black), unimodal (purple), or bimodal (red) distributions at each timepoint. Activity progresses from a null-distribution to unimodal and finally bimodal across-trial activity distributions, indicative of a bifurcation. B) Temporal generalization matrix. For stimuli at the detection threshold (about 50% detection rate), a classifier trained to decode trial outcome (hit/miss) from the activity pattern at each time point in a training data set is used to predict outcome based on the activity at each trial timepoint in held-out data. A diagonal pattern (e.g. in the lower-left dashed box) indicates a quickly-changing dynamical code. A square pattern (e.g. in the upper-right dashed box) indicates a stable code. C) Cortical surface representation of the mean and standard deviation of the normalized decoder coefficients for early (0-50ms) and late (300-350ms) periods of the trial. D) Mean (+,- SD) of the normalized decoder coefficients for early (0-50ms) and late (300-350ms) periods of the trial, as a function of the hierarchical position of each cortical area. E) Correlation (Pearson’s r) between the decoder coefficients at each timepoint and the cortical hierarchy. The red bar shows the time-range with a statistically significant correlation.

### A dynamic-to-sustained progression of activity states associated with ignition

Previous studies of conscious perception have reported that neural dynamics evolve from a dynamic to a relatively stable activity pattern (Dehaene and King 2016; Schurger et al. 2015). We aimed to decode the trial outcome (hit/miss) from activity at each timepoint for a fixed stimulus near to the threshold of detection using the temporal generalization method (King and Dehaene 2014; Meyers et al. 2008). We defined trial outcome based on activity in area 9/46d, and predicted this outcome using activity in all other cortical areas (therefore avoiding circularity). We observed a dynamic succession of patterns coding for stimulus visibility in the early trial stages, with reasonably high classification accuracy remaining close to the diagonal (Fig 3B, lower left box). In the later trial period we observed a stable pattern of high decoding accuracy, with the decoders trained between *∼* 100 *−* 400*ms* generalizing to all other timepoints within that range (Fig 3B, upper right box). Classifiers trained on some early timepoints had below-chance accuracy at decoding later timepoints (blue patches in Fig 3B). This indicates that early activity patterns are effectively reversed later in the trial. Similar results have been reported in the human experimental literature (King et al. 2016; Sergent et al. 2021). Our model suggests that the below-chance generalization from early to late timepoints may be due to higher associative areas sending net inhibitory feedback to areas that are lower in the visual hierarchy. Put another way, the stable, ignited activity pattern can lead to a reversal of the activity patterns that occur during stimulus propagation.

Decoding coefficients were highly variable over the first 50 ms (Fig 3C left, Fig 3D), before settling on a pattern (300-350ms) of high coefficients throughout a distributed network of frontal, parietal and some temporal regions. The standard deviation remains high only in regions of frontal cortex to which activity propagates last. The coefficients of the late decoder are higher in areas that are high in the cortical hierarchy (Fig 3D). A significant, and stable, correlation between decoder coefficients and the cortical hierarchy emerges late in the trial, after about 300ms (Fig 3E). This demonstrates how a stable code throughout fronto-parietal cortex can coexist with a dynamic activity in some areas of cortex (Fig 3C). This prediction can be tested experimentally.

### A dynamic bifurcation mechanism for ignition underlies stimulus detection

The previous analyses hinted at the possibility of a bifurcation occuring during stimulus detection. To better understand the dynamics determining whether individual trials would result in a hit or a miss, we built a simplified local model with a single area made of a single excitatory and a single inhibitory population (Fig. 4A). The equations are the same as in the full model, only the connectivity is different. Additionally, we focused on excitation mediated by the NMDA receptors. This reduces the system to two dynamical variables, corresponding to the synaptic variables *S*_*NMDA*_ and *S*_*GABA*_. This simplification enables us to analyze the dynamics of individual areas by looking at their phase portraits (Strogatz 2018).

**Figure 4:**
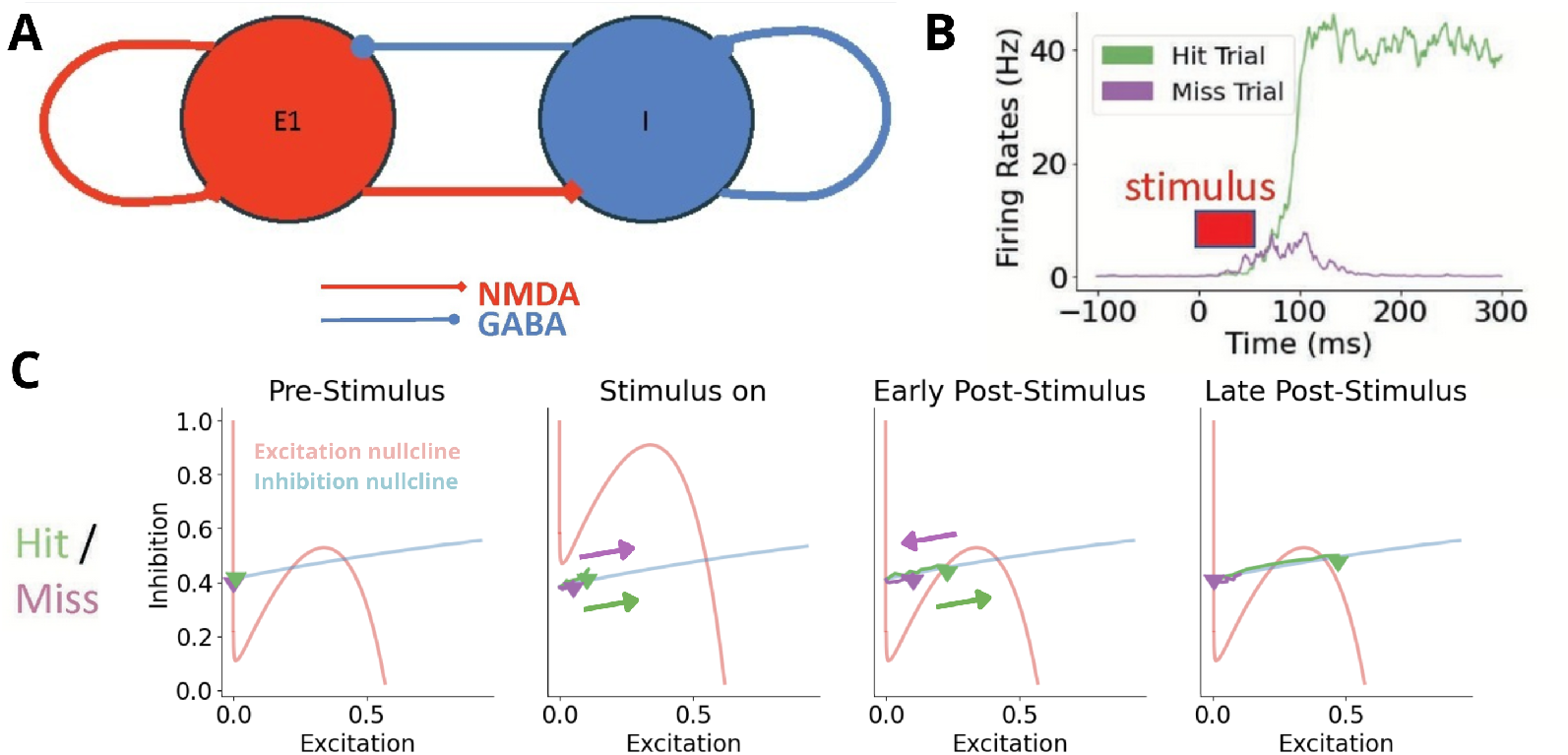
A dynamic bifurcation mechanism for ignition underlies stimulus detection. A) Simplified circuit for analysis of dynamics, containing a single excitatory and a single inhibitory population, interacting via NMDA and GABA receptors. B) Firing rates from area 9/46d in the large-scale system on individual hit (green) and miss (purple) trials. C) Phase portrait of the simplified network at different trial stages. Example dynamics for individual hit and miss trials are shown in green and purple, respectively. The stimulus causes the excitation nullcline (red) to move up, reducing the number of crossings with the inhibition nullcine (blue). Removal of the stimulus moves the nullclines back to the original positions. In the hit trial, by the time the stimulus has been removed, activity has reached the basin of attraction of the high activity steady state and progresses towards it (right nullcline crossing). In the miss trial, activity remains in the basin of attraction of the low activity steady state (left nullcline crossing), and returns towards it after removal of the stimulus..

We analyze the dynamics for hit and miss trials (Fig 4B, Supplementary Video 1). The system initially has two stable steady states (attractors), corresponding to low and high activity states (Excitation close to 0 and Excitation close to 0.5) and an unstable steady state (repeller, at about Excitation = 0.2). Before the stimulus, the system begins at the low steady state. The stimulus to the excitatory neural population shifts the excitation nullcline up, reducing the number of nullcline crossings from three to one. The single remaining crossing represents a stable steady state, and activity is attracted towards this high activity state during the stimulus. Due to noise, the speed at which the activity increases towards the nullcline crossing differs. When the stimulus is removed (’Early Post-Stimulus’), the nullclines rapidly shift back to their original position. As the unstable steady state (the middle nullcline crossing) repels activity away from itself, this effectively acts as a threshold. When the stimulus is removed, any activity to the left of the unstable steady state is attracted back to the low activity steady state, resulting in a miss, while any activity to the right is attracted towards the high activity steady state, leading to a hit (Fig. 4C). A stronger stimulus will lead to a larger shift in the excitatory nullcline, which increases the probability of trajectories reaching the basin of attraction of the high activity state (i.e. a hit). Therefore, the dynamic activity patterns of hits and misses can result from transient bifurcations induced by the external stimulus and noise.

### Fast propagation of stimulus information to prefrontal cortex depends on feedforward excitation mediated by AMPA receptors

The transient sensory activity and persistent prefrontal activity seen above and experimentally (Romo and Rossi-Pool 2020; Van Vugt et al. 2018) suggests that, unusually for a recurrent network, different parts of the cortex may act in different dynamical regimes (i.e. monostable vs bistable). Although the connectivity and spine count in our model are taken from anatomical data, the cell-type targets of the inter-areal connections (excitatory or inhibitory), and which glutamatergic receptors mediate these connections (AMPA or NMDA) are unknown. We therefore performed a parameter search to uncover the combinations of cell-type targets and glutamatergic receptors that can reproduce such dynamics.

We allowed the parameters for feedforward and feedback components of pathways to vary independently. We used the fraction of supragranular labelled neurons (SLN) as a validated marker of the degree of ’feedforwardness’ of a pathway (Markov et al. 2014), but did not explicitly include layers in the model. Here we refer to the sum of axons from any area to another as a ’pathway’, which is made up of a combination of SLN (i.e. pure feedforward) and 1-SLN (i.e. pure feedback) components. For example, the pathway from V1 to V2, which is a classic feedforward pathway, has an SLN of 0.72 (from the data in Froudist-Walsh et al. 2021).

We found three distinct regimes of model behavior in response to a brief, strong 50ms visual stimulus. Namely transient activity that returns to baseline (No Bistability), sustained activity across all cortical areas (Whole Cortex Bistability) and sustained activity only in association areas of cortex (Subnetwork Bistability) (Fig 5A,B). The reference parameter set used in all figures of the paper (unless specified otherwise) was taken from this Subnetwork Bistability regime.

**Figure 5:**
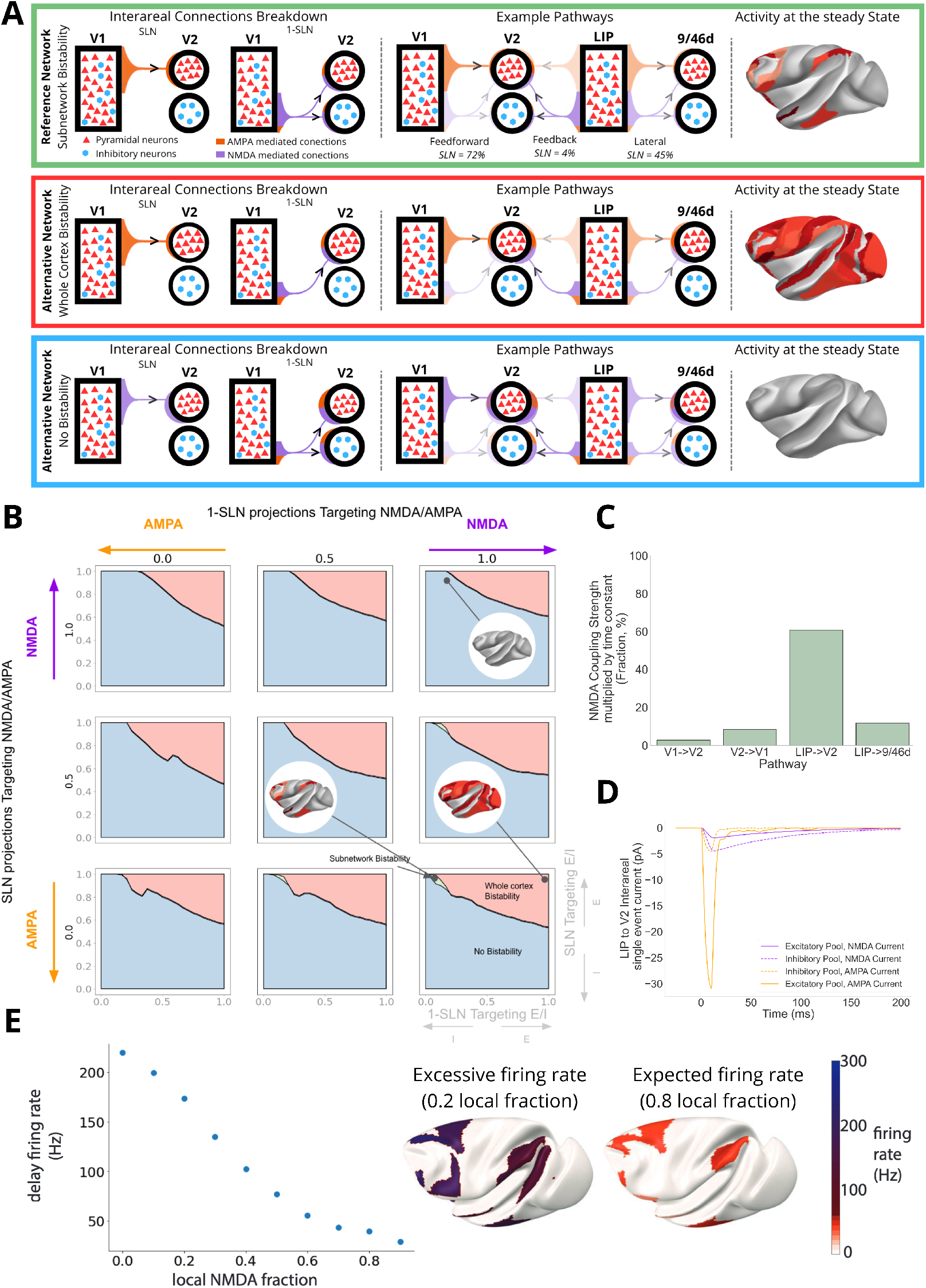
Rapid ignition depends on AMPA-dominated feedforward connections and balanced NMDA/AMPA-mediated excitation at feedback connections. A) Top. Representative connections of a parameter set resulting in subnetwork bistability. Left. Schematic of projections from V1 to V2. The target (Pyramidal vs Inhibitory neurons) and glutamatergic receptor (AMPA vs NMDA) mediating the connections depends on the SLN. Center. Three representative pathways, corresponding approximately to feedforward (V1 to V2, SLN = 0.72), feedback (LIP to V2, SLN = 0.04), and lateral pathways (LIP to 9/46d, SLN = 0.45). The opacity reflects the strength of each type of connection within the pathway. Right. The steady state firing rate across the cortex following a strong stimulus for this parameter set. Middle. Same as top, for a parameter set resulting in whole-cortex bistability. Bottom. Same as top, for a parameter set resulting in no bistability. B) Four-parameter search over glutamatergic receptors (AMPA/NMDA) and cellular targeting (excitatory/inhibitory) of inter-areal connections (performed separately for SLN and 1-SLN, corresponding approximately to feedforward and feedback components of each pathway). The models exhibit three distinct dynamic behaviors - subnetwork bistability (green), whole cortex bistability (red), and no bistability (blue) - with the former most consistent with empirical observations. Example parameter sets from Panel A are mapped onto this space for reference. C) Pathway specific NMDA contribution. Inter-areal NMDA coupling strength multiplied by the time constant - calculated as G_N_ τ_N_ /(G_N_ τ_N_ + G_A_τ_A_) - of the reference network for a feedforward pathway (V1 →V2), two feedback pathways (LIP → V2 and V2 → V1) and a lateral pathway (LIP → 9/46d). D) Inter-areal synaptic currents from LIP to V2 when all other inter-areal connections are removed and a brief stimulus is applied to LIP. E) Left. Average firing rate in areas showing delay activity for models with different local NMDA fractions. Right. Delay period activity across the cortex for models with different local NMDA fractions (Top: local NMDA fraction = 0.2, Bottom: local NMDA fraction = 0.8). Only the models with a relatively high fraction of local excitation mediated by NMDA receptors can reproduce realistic fronto-parietal activity levels.

We observed that to obtain subnetwork bistability dynamics, feedforward components of pathways in the model should target mainly excitatory cells, principally via AMPA receptors (Fig 5A-B). In contrast, the feedback components of pathways should target both excitatory and inhibitory cells, with a greater contribution of NMDA receptors.

We next performed a separate parameter search, to identify parameter sets that could replicate realistic propagation times (Figure S1). Notably, a strongly overlapping set of parameters was required to replicate rapid ignition within the realistic 130-200ms range (de Lafuente and Romo 2006; Thorpe and Fabre-Thorpe 2001; Van Vugt et al. 2018). Specifically, we found that feedforward components targeting excitatory cells via AMPA receptors and feedback components targeting inhibitory and excitatory cells, with a greater NMDA-dependent contribution allows the feedforward excitation to transiently ’escape’ the inhibition, and successfully propagate stimulus-related activity along the cortical hierarchy. Therefore, the pattern of feedforward AMPA-mediated excitation and mixed feedback (NMDA + AMPA) is required to replicate the spatiotemporal activity of sensory and associative areas during stimulus detection.

We next focused on one classic feedforward pathway (V1 → V2), two feedback pathways (V2 → V1 and LIP→ V2) and one lateral pathway (LIP → 9/46d, Fig 5C) from the cortex-wide model (reference parameter set). The NMDA fraction (expressed as 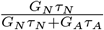 where *τ*_*N/A*_ is the receptor time constant) was highest in inter-areal feedback pathways, followed by inter-areal lateral pathways and lowest in feedforward pathways. However in all of these pathways there remains a significant AMPA contribution. We then cut all the inter-areal connections but LIP to V2 and injected a strong and brief (10ms) current in LIP. Despite LIP to V2 being a feedback connection, and having one of the largest NMDA contributions in the network, the peak of the resulting inter-areal AMPA current is over five times greater than the peak of the NMDA current (Fig 5D). Despite the dominance of AMPA at inter-areal connections overall, in our model it is an important observation that the NMDA coupling is stronger in feedback and lateral pathways than feedforward pathways, a prediction that remains to be measured experimentally.

### Ignition depends on NMDA receptor activation in local excitatory connections

Does ignition also depend on the receptors that mediate local intra-areal excitation? We adjusted the model so that the NMDA fraction of local excitatory connections varied. For a very low NMDA fraction, we see sustained activity in the fronto-parietal network, but at unrealistically high firing rates (mean frontoparietal delay activity = 173Hz for local NMDA fraction = 0.2, Fig 5E). Only models with a relatively high fraction of local excitatory connections mediated by NMDA receptors showed sub-network bistability and sustained activity in the fronto-parietal network at a reasonable firing rate (mean frontoparietal delay activity = 40Hz for local NMDA fraction = 0.8, Fig 5E). However, if local connections were completely mediated by NMDA receptors, bistability was lost. This suggests that a contribution of AMPA-mediated excitation is helpful to engage NMDA-mediated excitation and sustained activity. Therefore, our model suggests that NMDA receptors at local excitatory connections are crucial for ignition of cortical activity in response to a stimulus, in support of the previous theoretical (Wang 1999) and experimental (Wang et al. 2013) findings.

### The NMDA/AMPA ratio decreases along the cortical hierarchy

We calculated the NMDA fraction (as a fraction of NMDA and AMPA receptor-mediated excitation) for each area in the model (Subnetwork Bistability parameter set). In the model, we observed a strong decrease in the NMDA fraction along a 40-area cortical hierarchy (Fig 6A, *r* = −0.71, *p* = 3 × 10^*−*7^, hierarchy data from Froudist-Walsh et al. 2021). This decreasing NMDA fraction gradient is not seen in the networks that are incapable of producing realistic ignition dynamics (Whole-Cortex Bistability; No Bistability parameter sets) (Supplementary Figure 2). This leads to a testable non-trivial prediction: the cortical hierarchy, if capable of subnetwork bistability and rapid ignition, should have a decreasing NMDA fraction gradient.

**Figure 6:**
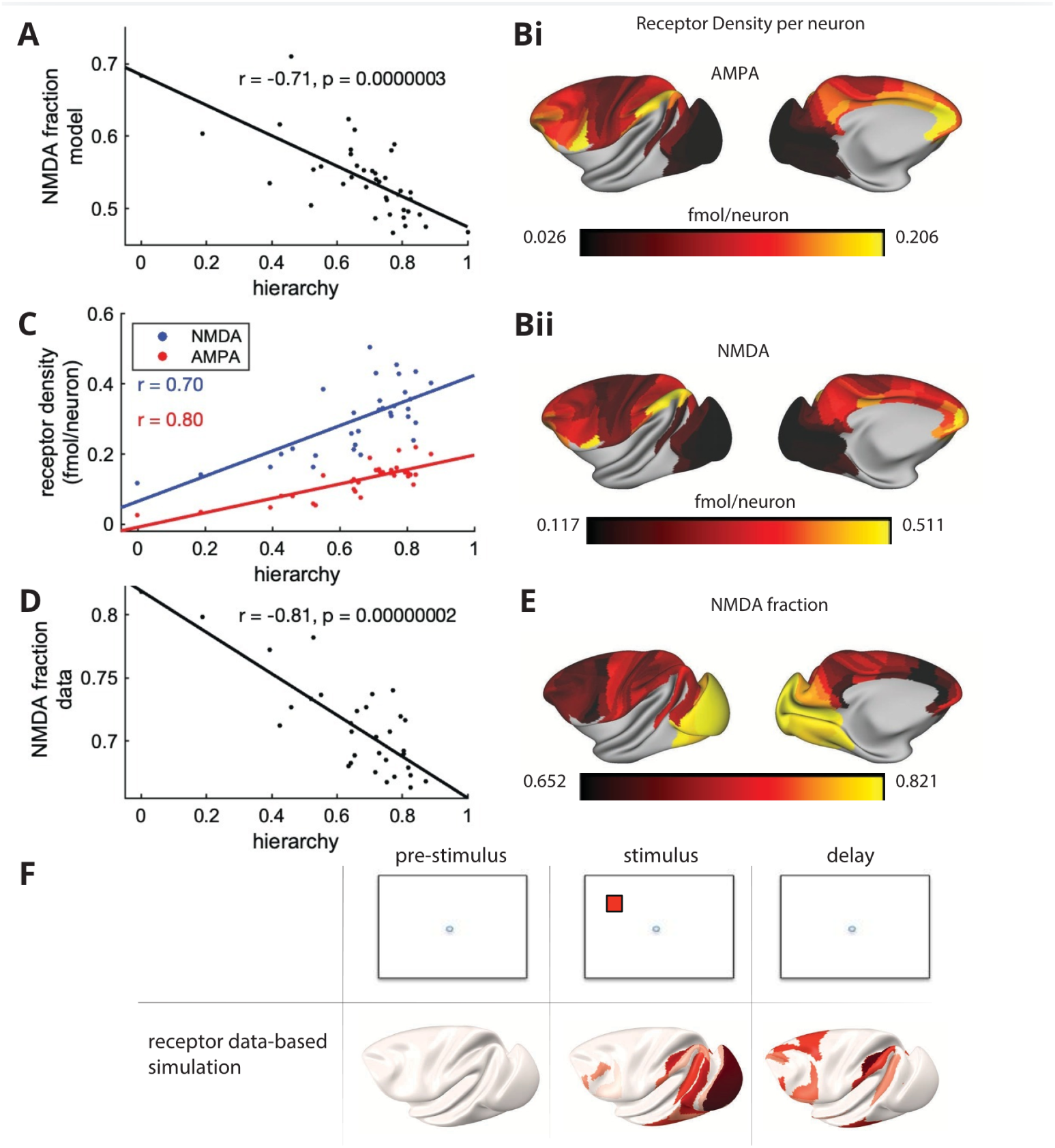
The NMDA/AMPA ratio decreases along the cortical hierarchy and supports ignition. A) The fraction of excitatory inputs via NMDA receptors (compared to total NMDA+AMPA inputs) in the model decreases along the hierarchy. B) The density of i) AMPA and ii) NMDA receptors across 109 regions of macaque cortex. Receptor density was measured using in-vitro receptor autoradiography, and divided by the neuron density data from Collins et al., 2010. C) AMPA and NMDA densities per neuron both increase along the cortical hierarchy. D, E) The fraction of NMDA receptors (compared to total NMDA+AMPA receptors) in the macaque receptor autoradiography data decreases along the hierarchy. F) The model was adjusted to match the receptor densities observed in the autoradiography data. The receptor data-based model shows ignition of fronto-parietal activity in response to a visual stimulus.

We tested this prediction by analyzing *in-vitro* receptor autoradiography data from 109 regions of macaque cortex (Froudist-Walsh et al. 2023; Impieri et al. 2019; Niu et al. 2020, 2021; Rapan et al. 2021). By dividing the receptor density by the neuron density in each area (Collins et al. 2010; Froudist-Walsh et al. 2021, 2023), we were able to estimate the NMDA and AMPA density per neuron in each area (Fig 6B). We found that both the NMDA (*r* = 0.70, *p* = 1 × 10^*−*5^) and the AMPA (*r* = 0.80, *p* = 3 × 10^*−*8^) receptor densities per neuron increased along the cortical hierarchy (Fig. 6C). We defined the ’NMDA fraction’ in each area as the NMDA receptor density divided by the sum of the NMDA and AMPA receptor densities. Despite the increases in both NMDA and AMPA densities along the hierarchy, there was a strong negative correlation between the NMDA fraction from the experimental data and the cortical hierarchy (Fig 6D,E, *r* = *−*0.81, *p* = 2 *×* 10^*−*8^), confirming our model prediction.

### The decreasing NMDA/AMPA gradient supports ignition

We then adjusted the model parameters to match the experimentally observed NMDA and AMPA receptor densities and spine counts (Niu et al. 2024). In Figure 5 we demonstrated how the NMDA fraction at feedforward and feedback connections critically determines ignition time. We therefore maintained the NMDA fraction at feedforward and feedback connections from the reference model (an assumption on proximity of this aspect of the reference model to biology). However, we allowed the NMDA fraction at local connections to vary across areas, so that the overall NMDA fraction of each area in the model closely matched that observed in the receptor autoradiography data. Without changing any other parameters, we observed that this receptor data-based simulation displayed subnetwork bistability dynamics (Fig 6F). Therefore, our previous result is robust to a more biologically-constrained parameter set. This supports our prediction that a decrease in the NMDA fraction along the hierarchy may have evolved to enable the dynamic inter-areal interactions required to support ignition-like dynamics.

## Discussion

We developed a novel large-scale model of monkey cortex and used it to simulate a detection task. The present work builds upon a previously proposed global neuronal workspace (GNW) architecture (Dehaene et al. 2003; Dehaene and Changeux 2005), but now incorporating newly available weighted and directed cortical connectivity data (Froudist-Walsh et al. 2021; Markov et al. 2014, 2012; Markov et al. 2013; Mejias et al. 2016) and receptor data (Froudist-Walsh et al. 2023; Impieri et al. 2019; Niu et al. 2020, 2021; Rapan et al. 2021). Using this model, we replicated multiple spatial and temporal features of neural activity that are observed during conscious perception. The close correspondence of our simulations to experimental results enabled us to identify candidate synaptic and network level mechanisms underlying conscious perception. Our model predicts that the rapid ignition of fronto-parietal activity that accompanies conscious perception depends on feedforward inter-areal excitation being primarily mediated by AMPA receptors. However, AMPA receptors are not sufficient to reproduce experimental activity patterns, as networks that rely on AMPA receptors for local excitation cannot reproduce realistic firing rates in fronto-parietal cortex. We show that sustained fronto-parietal activity depends critically on the NMDA receptors. Our modeling results led to the surprising prediction that the NMDA/AMPA ratio should decrease along the cortical hierarchy. We confirmed this model prediction by analyzing *in-vivo* receptor autoradiography data across dozens of cortical regions. Finally, we show how this decreasing NMDA/AMPA ratio supports the fast propagation of stimulus-related information along the hierarchy, and ignition of a distributed fronto-parietal network, as is seen during conscious perception in both human and nonhuman animals.

### Network, cellular, and synaptic mechanisms of conscious access

Our model displayed behavior and cortical activity patterns that fit well with several results from the experimental literature. Notably, early sensory cortex displayed a transient, approximately linear response to the stimulus. This contrasted with the late, sustained, all-or-none response in parts of the prefrontal cortex, reproducing the experimental findings in a monkey experiment specifically designed to test the GNW theory (Van Vugt et al. 2018). Our model suggests that similar transient signals should be detected throughout the early visual system, while an all-or-none response should depend on the fronto-parietal circuit. This all-or-none distributed activity broadly lends support to the GNW framework (Dehaene et al. 1998, 2003), in which a strongly interconnected network of prefrontal and parietal neurons broadcasts information widely throughout the brain. Some models of the GNW theory suggest that this fronto-parietal network would excite, sustain and broadcast the original stimulus representation in sensory neurons after the stimulus has disappeared (Dehaene et al. 2003). While such delayed activity in early sensory cortex has been observed in some studies (Harrison and Tong 2009; Supèr et al. 2001; van Kerkoerle et al. 2017), the consistency with which it occurs is not clear, and many studies have not observed sustained neural activity (Leavitt et al. 2017), or only a small modulation (e.g. Van Vugt et al. 2018). To reproduce these latter experimental activity patterns, we found that feedback connections must substantially target inhibitory neurons. This is consistent with recent reports that top-down attention has a net-inhibitory effect on sensory cortex (Huang et al. 2019; Javadzadeh and Hofer 2022; Yoo et al. 2021) and with previous models predicting the need for feedback inhibition to sustain distributed persistent activity (Mejias and Wang 2022). A parsimonious explanation of the contrasting data on early sensory activity is that many tasks can be performed by maintaining a representation of only higher-order aspects of the stimuli, and therefore sustained activity is not required in early sensory cortex. This frees early sensory cortex to encode new sensory information. However, a focal top-down excitatory connection onto excitatory neurons in the target sensory area may be engaged, for example when attention to precise sensory details is required. The mixed excitatory and inhibitory effects of top-down connections can be addressed in more detail in future large-scale models with a greater diversity of neural populations in each area, divided across layers. This would also enable fuller exploration of the effect of asymmetries in feedforward and feedback connections on realistic cortical circuits and enable deeper integration with anatomical data. Such models could shed light on how different cognitive states may affect the dynamics (Melloni et al. 2011) and activity patterns across cortical regions as well as subcortical structures.

Our model suggests that at the synaptic level, NMDA receptors are crucial for the ignition phenomenon. NMDA-dependent excitation is affected by various neuromodulators, which can enable (Yang et al. 2013), enhance (Arnsten et al. 2020; Galvin et al. 2020; Vijayraghavan et al. 2007), or counteract (Arnsten et al. 2020, 2019; Vijayraghavan et al. 2007) its effects. NMDA-spikes are relatively localized within pyramidal cell dendrites, but they increase the probability of plateau-like calcium spikes, which propagate a much greater distance towards the soma (Aru et al. 2020). This dendrite-soma coupling fades under anaesthesia (Suzuki and Larkum 2020). Moreover, activation of the apical dendrites of subcortically-projecting layer 5 pyramidal cells in the mouse somatosensory cortex is robustly associated with stimulus detection (Takahashi et al. 2020). Activation of these layer 5 pyramidal cells could trigger the subcortical vigilance signal required for ignition of the distributed cortical network (Müller et al. 2023; Munn et al. 2023; Whyte et al. 2024). Therefore, our finding of the importance of NMDA receptors and a (potentially subcortical) vigilance signal to ignition is consistent with multiple converging lines of evidence, which are illuminating the network, cellular and synaptic mechanisms of conscious processing.

### A dynamic-to-stable transition of cortical activity during conscious perception

Our model is agnostic as to whether the entire late sustained distributed activity pattern represents conscious access alone, or whether some of the activity could prepare a conscious report, or other post-access conscious processing (Cohen et al. 2020; Dellert et al. 2021; Pitts et al. 2018, 2012, 2014; Schlossmacher et al. 2020; Sergent et al. 2021). The task that we simulated is not able to distinguish activity related to conscious access and response preparation. Sergent and colleagues aimed to disambiguate these signals by presenting the same auditory stimuli to subjects during a task and during task-free listening (Sergent et al. 2021). They found that the same stimulus leads to late sustained activity on some trials, and not others. This late neural activity is predictive of both task-related reports, and reports of conscious contents that are randomly sampled during task-free listening. This bimodal distribution of late activity across trials is like what we see in the model. Late activity for both paradigms engaged a distributed fronto-parieto-temporal network. However, only the report paradigm recruited areas around the premotor and supplementary motor cortex. This suggests some separation of brain activity related to conscious-access and motor preparation. Simulation of activity patterns during cognitive tasks in connectome-based dynamical models is in its infancy, and the types of tasks that have been simulated are particularly simple (Ding et al. 2024; Froudist-Walsh et al. 2021; Mejias and Wang 2022; Zou et al. 2023). Future models may attempt to tackle more complex task structures and disambiguate patterns of activity related to conscious access and distinct post-access processes. This will require the development of large-scale models that are capable of selecting when to recruit motor programs.

The model in this paper bears some resemblance to recent large-scale models of working memory (Froudist-Walsh et al. 2021; Mejias and Wang 2022). While working memory signals generally persist for a period of a few seconds, conscious perception can be much shorter, on the order of 200ms. Our results suggest a shared mechanism between ignition and persistent activity associated with working memory. Behaviorally, conscious access can be brief and incessantly switch to different percepts or thoughts, whereas working memory storage is longer lasting. However, the difference between the two may not be fundamental. Indeed, a classifier trained to decode the visibility of a stimulus in a conscious perception task from MEG activity can generalise to decoding visibility from the early stages of activity during a working-memory delay period (Trübutschek et al. 2017). The differences in later activity patterns could be explained by how a sustained signal is turned off in the brain, such as by a corollary discharge signal during a motor response. This theoretical hypothesis remains to be tested experimentally. Other aspects of the neural dynamics that separate conscious from unconscious processing, including the complexity of pre- and post-stimulus neural activity (Goldman et al. 2019), and transient synchronisation (Melloni et al. 2007) were not investigated here. These types of dynamics may be related to attractors other than steady states, such as chaotic attractors and limit cycles (Wang 2021).

### Asymmetric feedforward and feedback excitation via AMPA and NMDA receptors reconciles contrasting anatomical and physiological findings

We show that ignition of distributed fronto-parietal activity can occur rapidly, when inter-areal feedforward excitation is predominantly mediated by AMPA receptors. In the model, when feedforward excitation was mediated by AMPA receptors, ignition occurred within about 200ms. This figure closely matches the timescale of ignition of prefrontal activity in monkeys performing a stimulus detection task (Van Vugt et al. 2018), and is similar to (but slightly quicker than) proposed neural signals of conscious perception in the larger human brain (Del Cul et al. 2007; Salti et al. 2015; Sergent et al. 2005; Sergent et al. 2021). Realistically fast access of stimuli to consciousness therefore necessitates a relative increase of AMPA receptors in cortical areas that receive feedforward connections from the visual system.

The differences between feedforward and feedback connections are functionally important, but also quite small in quantitative terms. It is important to note that the asymmetry between feedforward and feedback connections that we highlight results from the analysis of a parameter search. This combination of parameters produces the dynamical activity patterns that best resemble those seen experimentally. Many studies have reported NMDA/AMPA ratios at excitatory synapses (Feldmeyer et al. 1999; Gil and Amitai 2000; Groc et al. 2002; Jones and Baughman 1988; Kumar and Huguenard 2001; Markram et al. 1997a; McBain and Dingledine 1992; Myme et al. 2003; Umemiya et al. 1999; Watt et al. 2000), though they have not separated interareal feedforward, feedback and local recurrent connections. These studies invariably find both NMDA- and AMPA-mediated contributions to excitatory synaptic transmission, with the ratio depending on a variety of experimental details. However none of the above studies report either NMDA or AMPA dominating excitatory synaptic transmission at relatively high membrane potentials. This suggests that, in total, we should not expect purely NMDA nor purely AMPA dominated excitatory transmission. Only limited experimental evidence exists regarding the NMDA fractions at feedforward and feedback connections (Self et al. 2012) with this study providing indirect evidence for a role of AMPA receptors in feedforward visual information propagation, and for NMDA receptors in slower responses driven by recurrent connections (either local or inter-areal feedback). Similarly, few papers examine the net excitatory or inhibitory effects of feedforward and feedback connections (Huang et al. 2019; Javadzadeh and Hofer 2022; Yoo et al. 2021). However, one recent paper shows that feedback connections have a slight net inhibitory effect on their targets, which contrasts with the strong net excitatory effects of feedforward connections (Javadzadeh and Hofer 2022). In the future, these model predictions should be tested experimentally, including with ultrastructural analyses. In our model, the local inhibitory population was driven by NMDA-rich excitatory connections. The NMDA-mediated excitation is reminiscent of somatostatin-expressing (SST+) inhibitory neurons (Morabito et al. 2024). The SST+ cells are also major targets of feedback connections in the mouse visual system (Shen et al. 2022). Together, this indicates a potential role for SST+ neurons in the circuit. However, in the future AMPA input to inhibitory neurons and a diversity of cell-types should be included.

Physiological and computational studies of recurrent excitation in local circuits have emphasized the role of AMPA receptors in primary visual cortex (Yang et al. 2018) and NMDA receptors in dlPFC (Wang et al. 2013; Wang 1999). Reliance on the fast-acting AMPA receptors may facilitate the rapid encoding of stimulus information in V1, while the slow dynamics of the NMDA receptor are thought to be crucial for the characteristic persistent activity seen during working memory tasks. By going beyond local circuit considerations, we showed in this work that the density of both NMDA and AMPA receptors per neuron increases along the cortical hierarchy, while the NMDA/(NMDA+AMPA) fraction decreases. The decrease in the NMDA fraction along the hierarchy is in agreement with our modeling prediction and recent reports in the human brain (Goulas et al. 2021; Zilles and Palomero-Gallagher 2017). The increasing density of NMDA receptors along the hierarchy, may account for the commonly observed persistent activity in dlPFC, that is largely absent in V1 (Leavitt et al. 2017). This parallels the increases in dendritic spines on layer 3 pyramidal cells, which is the most common site of local recurrent excitatory connections (Elston 2007; González-Burgos et al. 2019). We show that NMDA at local connections are crucial for sustained activity to emerge. While much work has contrasted the response of V1 neurons during stimulus presentation to dlPFC neuron activity during the delay period of working memory tasks, some neurons in dlPFC also react transiently to a stimulus, although with a delayed onset. The transient response of these ’Cue’ neurons in dlPFC is greatly reduced by AMPA receptor antagonists (Wang et al. 2013; Yang et al. 2018). This is consistent with feedforward excitation of ’Cue’ neurons in dlPFC being mediated by AMPA receptors. Persistent activity in contrast is more reliant on NMDA receptors. However, NMDA receptors cannot act on their own in this function, as excitation via AMPA or nicotinic receptors is also required to depolarize the cell and remove the magnesium block to engage NMDA receptors (Van Vugt et al. 2020; Yang et al. 2013). NMDA receptors in V1 are largely responsible for modulating late responses via inter-areal feedback connections, rather than the initial feedforward pass (Self et al. 2012). As all the inter-areal cortico-cortical connections that V1 receives are essentially feedback, this may account for the larger NMDA/AMPA ratio in V1, even if relatively few local recurrent connections are via NMDA receptors. Our simulations show that a model that closely matches the experimentally-observed NMDA and AMPA distributions could produce robust stimulus propagation and ignition of fronto-parietal activity. The receptor-based model produced ignition-like dynamics without the need for adjustment to any parameters in the model. This is because the requirement for a decreasing NMDA/AMPA ratio can be naturally fulfilled with feedforward excitation mediated by AMPA receptors, and a greater contribution of NMDA receptors to feedback connections. We found that, in the model, a dominant AMPA-mediated excitation at local recurrent connections led to unrealistically-high firing rates. This leads to the following prediction. The relative contribution NMDA-mediated excitation increases from interareal feedforward connection, to interareal feedback connections, to local recurrent connections. To our knowledge, this has not yet been explicitly measured, and is a testable prediction of the model.

### Integration of the model in the consciousness literature

The present work proposes a neural mechanism for conscious access, the cognitive function that lets a stimulus enter in the current stream of consciousness (James 1890) and makes it reportable, verbally or non-verbally (Baars 2005; Dehaene et al. 1998). The other cognitive functions associated with consciousness, such as metacognition, self-awareness, or any form of attention, are not addressed. Two major current theories of consciousness, among many (Seth 2007), are Global Neuronal Workspace theory (Dehaene et al. 2003) and Integrated Information Theory (IIT) (Tononi 2004). Our model fits in the GNW literature, as it possesses the major characteristics of the Global Workspace, namely independent sensory modules competing to pass their information to a widely distributed set of areas that broadcast the information to vast parts of the cortex. It differs from previous computational models of the Global Neuronal Workspace in that it is built explicitly on mesoscopic connectome data, and therefore makes predictions for cortex-wide neural activity during detection tasks, as well as receptor distributions across the cortex. The associative areas responsible for ignition in the model, heterogeneously connected by the FLN matrix, resemble the core of the mesoscopic connectome (Markov et al. 2013) as well as the specialized majority network taken as an example of a high Phi complex (Oizumi et al. 2014) as stated by the IIT for the origin of Phenomenological Consciousness (Tononi 2004). However the location of the cortical areas, predominantly in frontal and parietal cortex seems to fit more precisely with GNW theory than IIT, which attributes conscious perception primarily to a posterior cortical ”hot zone” (Koch et al. 2016). Further anatomically-constrained large-scale modelling, or analysis of our model, could make explicit the areas of agreement and disagreement between GNW and IIT. Future work addressing aspects of metacognition, attention, predictive processing, emotional awareness, conscious volition or conscious thinking could aim to further bridge GNW with other prominent theories of consciousness (Fleming and Lau 2014; Graziano and Webb 2015; He 2023; Lau and Rosenthal 2011; Seth and Hohwy 2021; Shea and Frith 2019), and provide much-needed testable predictions about behaviour and neural dynamics to distinguish between such theories (Demertzi et al. 2019; Melloni et al. 2021; Yaron et al. 2021).

Our current model was specifically designed to replicate brain activity triggered by a brief, faint stimulus. This approach is often used in the study of neural correlates of conscious perception (Sergent et al. 2021; Van Vugt et al. 2018). Despite this, there is a host of other experimental paradigms where the identical stimulus can alternatively be perceived or missed. These paradigms include masking (Del Cul et al. 2009; Kouider and Dehaene 2007), attentional blink (Asplund et al. 2014; Kranczioch et al. 2005), and binocular rivalry (Hesse and Tsao 2020; Kapoor et al. 2022). Recent advancements in no-report binocular rivalry experiments have unveiled precise patterns of activity in temporal and prefrontal cortex. These studies allow for effective decodability of the conscious percept in both the Infero-temporal cortex (Hesse and Tsao 2020) and the lateral prefrontal cortex (Kapoor et al. 2022). Future work could investigate the network activity during such experiments, replicating these precise electrophysiological results and enabling predictions at the whole cortex scale.

According to the Global Workspace Theory, this sudden ignition of neural activity is the neural correlate of the broadcasting of information across the brain, which corresponds to the moment a stimulus reaches conscious awareness (Conscious Access). By proposing a specific neurobiological implementation of this hypothesis, our model makes predictions at the theoretical and mechanistic levels. The theoretical prediction is that a dynamic bifurcation, spatially distributed across the brain underlies the ignition phenomenon. The mechanistic prediction is that, to support the ignition phenomenon at realistic timescales, NMDA-mediated excitation must be i) low at inter-areal feedforward synapses, ii) relatively higher at inter-areal feedback synapses, iii) relatively high at local recurrent synapses. Global Workspace Theory can be invalidated by demonstrating that ignition does not correspond to the access of a stimulus to conscious awareness (e.g. Conscious Access without Ignition or vice versa). However, note that even if GNW happens to be wrong, and the ignition event recorded in the previous experiments is not a signature of conscious access, our work still provides a mechanistic hypothesis for these large scale cortical activity patterns (while no longer describing conscious access). It is possible to disprove our theoretical prediction by uncovering an alternative dynamical mechanism for ignition. It is also possible to disprove our mechanistic hypothesis, by showing that ignition relies on a distinct neurobiological mechanism. Experimentalists can also disprove our mechanistic hypothesis by measuring NMDA fractions at inter-areal (feedforward, feedback) and local recurrent connections that are inconsistent with our prediction. Therefore, it is possible to disprove the model, and to disprove GNW. However, disproving the mechanistic hypothesis of the model is not sufficient to disprove GNW.

## Conclusions

We built a connectome-based dynamical model of primate cortex, that successfully accounted for salient results on the spatiotemporal activity and behavior of primates performing tasks designed to assess conscious access. Our model predicts that feedforward excitatory connections should be dominated by AMPA receptors for rapid propagation of stimulus-related activity, while NMDA receptors in local recurrent connections and feedback projections are required for the ignition and sustained activity that accompanies conscious access. Our model reconciles seemingly contradictory anatomical and physiological data on the relative proportion of AMPA and NMDA receptors along the cortical hierarchy, and takes a step towards a cross-level (bridging network, cellular and synaptic mechanisms) theory of consciousness.

## Acknowledgements

This work was funded by the NIH grant R01MH062349, ONR grant N00014, NSF NeuroNex grant 2015276, ONR grant N00014, James Simons Foundation grant 543057SPI, the Swartz Foundation to XJW; UKRI BBSRC grant BB/X013243/1 to SFW, NIH/NSF CRCNS grant R01MH122024 to XJW and NPG. SD was supported by Collège de France, INSERM and CEA. NPG, LR and MN were supported by the Federal Ministry of Education and Research (BMBF) under project number 01GQ1902. We thank Guillermo González-Burgos for comments on an earlier draft of the manuscript.

## Materials and Methods

### Model overview

We developed a connectome-based dynamical model of the macaque cortex to investigate the synaptic and network mechanisms underlying the ignition of distributed neural activity that accompanies conscious perception. We simulated local cortical circuits at each of 40 cortical areas and set the existence and strength of directed connections between areas using retrograde tract-tracing data. Cortical areas differed based on their inter-areal connectivity and dendritic spine count on pyramidal cells. As a starting point, we adapted a recently developed model of distributed working memory in 30 cortical areas (Mejias and Wang 2022).

### Retrograde tract-tracing data

The inter-areal connectivity data in this paper was acquired by Henry Kennedy and colleagues as part of an ongoing effort to map the cortical mesoscopic connectome of the macaque using retrograde tract-tracing (Markov et al. 2014, 2012; Markov et al. 2013; Mejias et al. 2016). Here we use the directed, weighted connectivity data between 40 cortical areas, which is the most recent release (Froudist-Walsh et al. 2021).

A few details of how the connectivity data was collected and processed will help the reader understand the connectome-based dynamical model. For each target area, a retrograde tracer was injected into the cortex. The tracer was taken up in the axon terminals in this area, and retrogradely transported to the cell bodies of neurons that projected to the target. The cortical areas (*l*) that send axons to the target area (*k*) are called source areas. For a given injection, all marked cell bodies in the cortex outside of the injected (target) area was counted as labeled neurons. The number of labeled neurons (*LN*) within a source cortical area was then divided by the number of labeled neurons in the whole cortex (excluding the target area), to give a fraction of labeled neurons (FLN). The FLN was averaged across all injections in a given target area. For this calculation, we include all cortical areas (*n*^*areas*^ = 91) defined in the Lyon atlas (Markov et al. 2012).

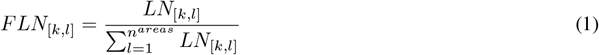

Note that there are 91 cortical areas in the Lyon atlas, and currently 40 areas have been injected with retrograde tracers. This gives the connection strength from all 91 areas to the 40 injected areas, and the full bidirectional connectivity of a subgraph of 40 areas. We use this 40-area subgraph as an anatomical basis for the dynamical model.

In addition, for each inter-areal connection we defined the supragranular labeled neurons (SLN) as the fraction of neurons in the source area whose cell bodies were in the superficial (aka supragranular) layers.

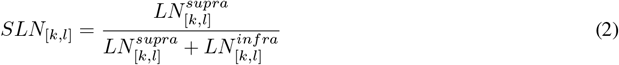

The subiculum (SUB) and piriform cortex (PIR) have a qualitatively different laminar structure to the neocortical areas, and therefore supra- and infra-laminar connections (and thus the SLN) from these areas are undefined. We removed all connections from these areas from the following calculations (*n*^*areas,SLN*^ = 89). These connectivity data are available on the core-nets website (register, click the “Download” button, and select the data associated with Froudist-Walsh et al. 2021).

### Dendritic spine data

The spine count data were taken from a series of studies by Elston and colleagues (Elston 2007) and mapped onto the Yerkes19 cortical surface (Donahue et al. 2016), as described in (Froudist-Walsh et al. 2021, 2023). Locations on the Yerkes19 cortical surface are represented by 32,492 vertices. The spine count data was obtained by Elston and colleagues from 27 injection sites across the cortex. For each injection site we estimated the number of vertices overlapping with each area in the Lyon atlas. If a cortical area contained only one injection site, the mean spine count from pyramidal cells in that site was taken as the spine count for the area. If a cortical area contained multiple injection sites, we performed a weighted average of the spine counts, according to the number of vertices of overlap. In this way we estimated the spine counts on pyramidal cells in 24 of the 40 injected regions in the Lyon atlas. Based on the strong positive correlation between spine count and cortical hierarchy (r = 0.61, p = 0.001), and following previous work (Chaudhuri et al. 2015; Froudist-Walsh et al. 2021; Mejias and Wang 2022), we inferred the spine count for the remaining regions based on the hierarchy using linear regression.

### Local cortical circuit architecture

In each cortical area we simulated a local circuit, with two interacting excitatory populations (*E*_1_ and *E*_2_), and one population of inhibitory (*I*) neurons. This is based on a mean-field reduction of a spiking neural network model of cortex (Wang 2002; Wong and Wang 2006).

### Description of dynamical variables

The neural populations interact via synapses that contain NMDA, AMPA and GABA receptors. Each receptor has its own dynamics, governed by the following equations.

The synaptic variables are updated as follows (Wang 1999; Wong and Wang 2006)

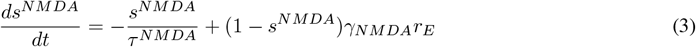

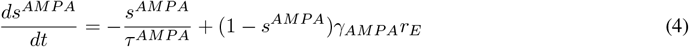

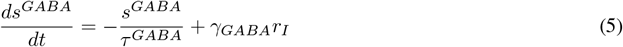

where *s* is the fraction of open synaptic ion channels due to bound receptors, *τ* is the time constant of decay of that receptor and *γ*_*NMDA*_, *γ*_*AMP A*_ and *γ*_*GABA*_ are constants. *r*_*E*_ and *r*_*I*_ are the firing rates of the presynaptic excitatory and inhibitory cells that stimulate the NMDA, AMPA and GABA receptors, calculated below.

### NMDA/AMPA ratio

We explored the effects of different NMDA/(NMDA+AMPA) fractions, *κ*, at local and long-range feedforward and feedback connections. The values used for the main simulations, unless otherwise stated, are in Table 2.

### Modulation of excitatory connections by dendritic spines

Approximately 90% of excitatory synapses on neocortical pyramidal cells are on dendritic spines (Nimchinsky et al. 2002). On this basis, we modulate the strength of excitatory connections according to the dendritic spine count.

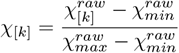

for all cortical areas 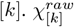 is the spine count for area *k*, and 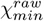 and 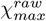 are the minimum and maximum spine counts observed in the data. *χ*_[*k*]_ is therefore the spine count of area *k* rescaled to lie in the [0, 1] range.

We then apply the gradient of excitation as follows.

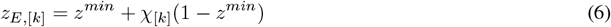

where *z*^*min*^ sets the lower bound for the modulation of excitatory connections by the spine count, *χ. z*_*E*,[*k*]_ therefore defines how spine count modulates excitatory connections in area *k*.

### Description of local currents

The local NMDA current onto each population *Ei* ∈ *{E*_1_, *E*_2_*}* in area [*k*] is calculated as follows

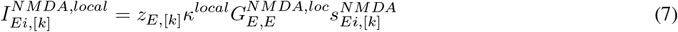

Where *z*_*E*,[*k*]_ is the dendritic spine count gradient, *κ*^*local*^ the NMDA receptor fraction of the postsynaptic population, 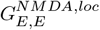 the local NMDA coupling from the population to itself.

Local connections tend to target the perisomatic area (soma and proximal dendrites) of pyramidal cells (Kalisman et al. 2005; Markram et al. 1997b; Petreanu et al. 2009). The soma and proximal dendrites act as a single functional compartment that is separate from a distal dendritic compartment (Yuste et al. 1994). As our dendritic function *F* (described below) models this distal dendritic compartment, we do not pass local excitatory connections through *F*.

Similarly local excitatory connections via the AMPA receptors are scaled by the AMPA receptor fraction 1 − *κ*^*local*^, the dendritic spine count gradient *z*_*E*,[*k*]_, and 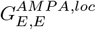 the local AMPA coupling from the population to itself.

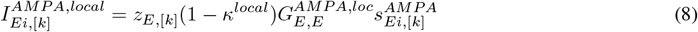

Local inhibitory connections are not directly modulated by the dendritic spine count (as spines indicate excitatory synapses on pyramidal cells, Nimchinsky et al. 2002).

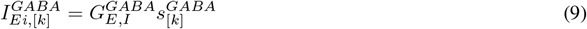

Where 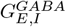 is the connection strength from the inhibitory pool to the excitatory pools.

In order to keep the spontaneous activity level similar across brain areas, the local NMDA input to the I population increases with the spine count, and is defined as followed (Mejias and ng 2022)

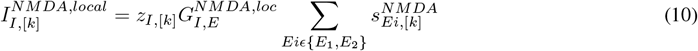

with

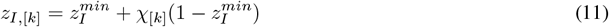

For the Main Figures in the manuscript, there is no local AMPA current targeting the inhibitory population. However, including AMPA input to inhibitory cells does not significantly change the results.

### Description of noise, background and vigilance currents

Noise is modeled as an Ornstein-Uhlenbeck process, separately for each population i in E1,E2,I.

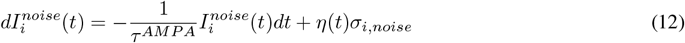

where *σ*_*i,noise*_ is the standard deviation of the noise and *η* is Gaussian white noise with zero mean and unit variance.

A constant background current 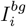 was also added to each population (Table 2). This represents input from brain areas that are not explicitly modeled.

In addition, we examined the effect of an extra, weak excitatory current, *I*^*vig*^, to each unit in associative areas (top 75% of areas ranked according to the hierarchy), which simulated the effect of vigilance on the model (Dehaene et al. 2003; Dehaene and Changeux 2005; Müller et al. 2023; Munn et al. 2023; Whyte et al. 2024).

As a simplification, each of these currents targets the perisomatic compartment (i.e, it is not passed through the distal dendritic function *F*).

### Large-scale connectivity structure

In the model, cortical areas are connected using connectivity strengths derived from the retrograde tract-tracing data. The long-range connectivity matrices are built from the FLN matrix. However, as noted in (Markov et al. 2014, 2012; Mejias et al. 2016), the FLN matrix spans 5 orders of magnitude. The relationship between anatomical and physiological connectivity strengths is not clear, but if we were to use the raw FLN values in the large-scale model, many of the weaker connections would become irrelevant. To deal with this, we follow Mejias et al. 2016; Mejias and Wang 2022 and rescale the FLN matrix in order to increase the influence of smaller connections while maintaining the topological structure. Mejias et al. 2016 found this rescaling was necessary to reproduce the significant inter-areal interactions found in (Bastos et al. 2015), and give a range of effective connectivity values similar to previous estimates (Song et al. 2005).

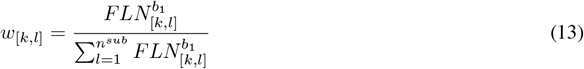

Here we restrict calculations to the injected cortical areas *i, j*, which allows us to simulate the complete bidirectional connectivity structure within the subgraph (*n*^*sub*^ = 40). Note that intra-areal connections are not quantified in the dataset. We use the same parameter value *b*1 as in (Mejias et al. 2016; Mejias and Wang 2022) (Table 2) to construct our inter-areal connectivity matrix *W*.

### Calculation of long-range currents

Excitatory cells in different cortical areas with the same receptive fields are more likely to be functionally connected (Zandvakili and Kohn 2015). This is reflected in our model as follows. In the source area, there are two excitatory populations, 1 and 2, each sensitive to a particular feature of a visual stimulus (such as a location in the visual field). Likewise in the target area, there are two populations 1 and 2, sensitive to the same visual features. We assume that the output of population 1 in the source area goes to population 1 in the target area, and the output of population 2 in the source area goes to population 2 in the target area.

The total long-range connections mediated by the NMDA receptors on the excitatory population *E*_*i*_ in area [*k*] is calculated as follows:

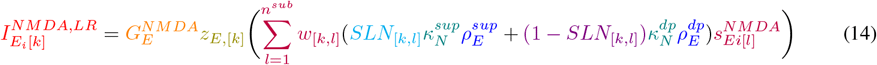

The amount of long-range current onto the excitatory population *E*_*i*_ in area *k* that comes via NMDA receptors depends on the fraction of open synaptic ion channels due to occupied NMDA receptors and the anatomical strength of inter-areal connections from all source areas *l* that target area *k*. This is scaled by the global excitatory NMDA coupling strength and the amount of dendritic spines per pyramidal cell (i.e. excitatory synapses) in area *k*. We separate the superficial layer from the deep layer projections as they may be mediated by different receptor types and target different cell types. Equations (16 - 19) can be understood similarly.

Note that distinct layers were not explicitly simulated in the model. However, in the real brain, connections from superficial and deep layers may have different impacts on brain dynamics. Therefore, we allowed the cell-type targets and receptors mediating interareal connections from superficial and deep layers to be variables. We investigate the impact of modifying these variables in Figure 2.

To be precise, 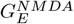 is the global coupling for NMDA-mediated inter-areal connections, *z*_*E*,[*k*]_ is the dendritic spine count value for area *k* (as defined above), *w*_[*k,l*]_ is the anatomical connection strength from area *l* to area *k, SLN*_[*k,l*]_ is the fraction of neurons projecting from area *l* to area *k* that have their cell bodies in the superficial layers (as above), 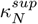 is the fraction of excitation that is mediated by NMDA receptors for connections from superficial layers, 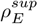 is the fraction of superficial layer projections targeting excitatory cells and 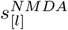 is the NMDA synaptic gating variable from the corresponding excitatory population in source area *l*. Similarly, 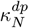 is the fraction of excitation from deep layer projections mediated by NMDA receptors, and is the fraction of deep layer 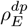 projections targeting excitatory cells. *i* = 1, 2 denotes the excitatory population.

Long-distance connections tend to target more distal parts of the dendrites (Petreanu et al. 2009), which act as a functionally separate compartment from the perisomatic area (Yuste et al. 1994). For this reason, we pass the long-distance connections through the dendritic function *F* before they reach the soma.

Similarly, the total long-range connections of the excitatory population in area [*k*] mediated by AMPA receptors is calculated as follows:

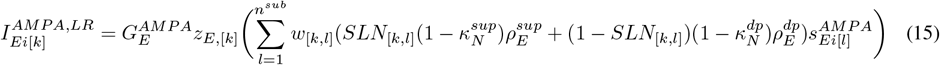

where 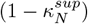 and 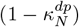 are the fraction of inter-areal connections from superficial and deep layers mediated by *AMPA* receptors. This is scaled by the global excitatory AMPA coupling strength 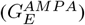.

The total excitatory long-range current in then computed as follow:

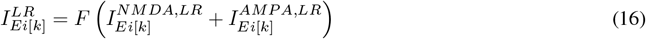

The function *F* is a simplification of a dendritic function used in previous local and large-scale models (Froudist-Walsh et al. 2021; Yang et al. 2016). It helps the network stabilize, and avoid epileptic behaviours.

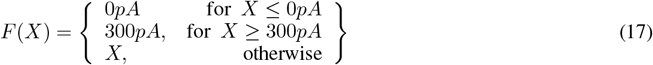

The total long-range connections targeting inhibitory population in area [*k*] that are mediated by NMDA receptors is calculated as follows:

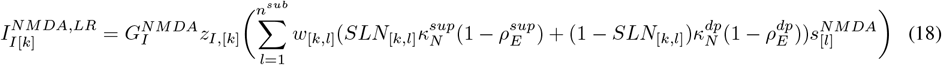

where 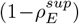 and 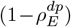 are the fraction of feedforward and feedback inter-areal connections targeting inhibitory cell populations. We assume different effective strengths for long-range connections targeting excitatory and inhibitory pools, captured by 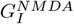 and 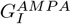. Although cortical inhibitory interneurons do not contain dendritic spines, we assume that the level of excitation onto inhibitory scales similarly with the spine count. This has been shown to be an effective way of maintaining spontaneous activity levels across areas (Mejias and Wang 2022).

The total long-range connections targeting the inhibitory population in area [*k*] that are mediated by AMPA receptors is calculated as:

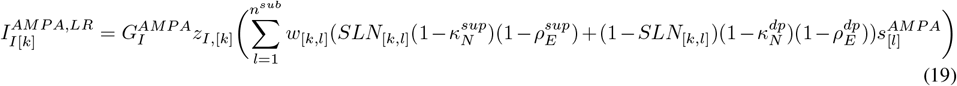

with all variables as described above.

### Application of external stimuli for tasks

In all simulations, the stimulus is applied for 50ms to excitatory population 1 in area V1. In the brain, visual input from LGN to V1 targets layer IV local excitatory neurons, which then excite the perisomatic areas of layer III pyramidal cells. For this reason we model external input to the perisomatic compartment of excitatory neurons in V1 (i.e., it is not passed through the dendritic function *F*). In all equations, the stimulus is designated by the term *I*^*stim*^.

### Total current in large-scale model

The total current for each neural population *i* in each area *k* equals the sum of all long-range, local and external inputs, and intrinsic currents,

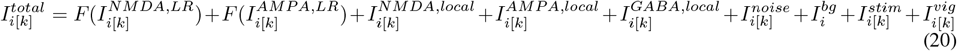

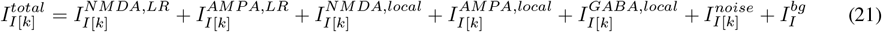

where *iϵ{E*_1_, *E*_2_*}*, (E1: excitatory population 1; E2: excitatory population 2; I: inhibitory population).

### Description of f-I curves

The f-I (current to frequency) curve of the excitatory population is

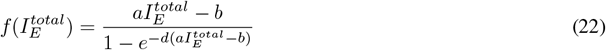

where 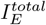 is the total input to the population, *a* is a gain factor, *d* determines the curvature of 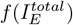, such that if *d* is large, 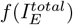 acts like a threshold-linear function, with threshold *b* (Abbott and Chance 2005).

The f-I curves for the inhibitory neuron populations are modeled using a threshold-linear function

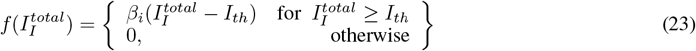

where 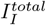 is the total input to the population, *β*_*i*_ is the gain and *I*_*th*_ is the threshold.

See Table 2 for parameter values.

The firing rates are updated as follows

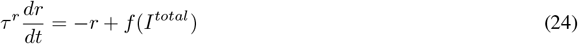

for all cell types.

### Classification of model dynamics

This corresponds to the analysis in Figure 3A.

### Model 0: Null Model

In this model, the external input has no effect on the activity. Irrespective of whether a stimulus was presented or not, and irrespective of its strength, activity follows a Gaussian distribution centered on *µ* with a standard deviation of *σ*.

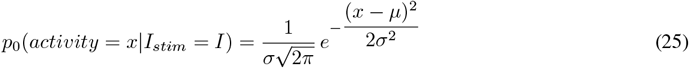

There are two free parameters: *µ* and *σ*

### Model 1: Unimodal Non-Linear

In this model, the activity evoked for each stimulus strength *I* follows a Gaussian distribution centered on a mean *µ* - following a sigmoid function of *I* - and a standard deviation *σ*, following a linear function of the mean *µ*. For this model, the probability to reach activity level *x* for an input *I* is given by:

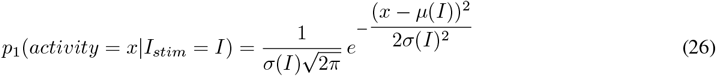

with

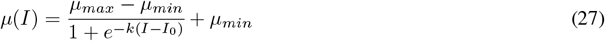

and

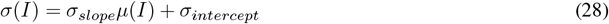

There are six free parameters: *µ*_*max*_, *µ*_*min*_, *k, I*_0_, *σ*_*slope*_ and *σ*_*intercept*_

### Model 2: bifurcation

In this model, the activity evoked for each stimulus strength *I* has a probability *β*(*I*) to belong to a high state (Gaussian distribution centered on *µ*_*high*_ of variance *σ*_*high*_), and a probability (1 *− β*(*I*)) to belong to a low state (Gaussian distribution centered on *µ*_*low*_ of variance *σ*_*low*_, the baseline activity observed in the absence of stimulation). For this model, the probability to reach activity level *x* for an input *I* is given by:

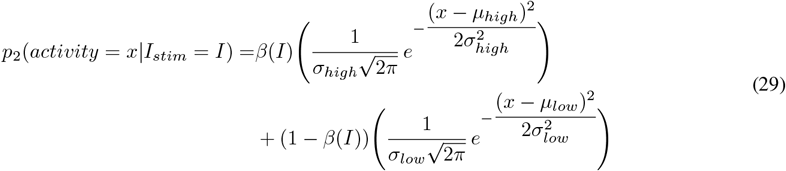

with

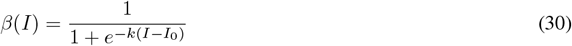

There are six free parameters: *µ*_*high*_, *µ*_*low*_, *σ*_*high*_, *σ*_*low*_, *k* and *I*_0_

### Bayesian model comparison

We compared the performance of the different models simulating 100 simulations for four different input current values (400 trials in total). The activity was sampled every 40ms and the activity was averaged over all 40 areas. The best parameters for each model were estimated by maximum likelihood, i.e., by finding the parameters maximizing the product of the likelihoods across the different trials (or, equivalently, maximizing the sum of the log likelihoods). The parameter search was achieved using the scipy.optimize function. In order to compare our different models, we used the following formula, where *P* (*M*_*i*_|*x*(*t*)) is the posterior probability of the model *i* ∈ {0, 1, 2} at the time step *t* (Lebarbier and Mary-Huard 2004).

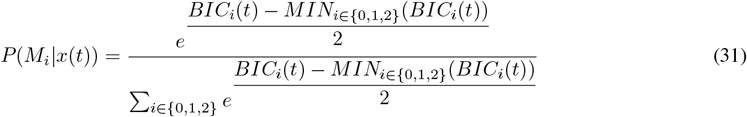

where *BIC*_*i*_(*t*) correspond to the Bayesian Information Criterion of model *i* at time step *t* for the best parameter set of this model.

### Temporal generalization of stimulus detection decoders

To decode the trial outcome from instantaneous trial activity patterns, we first separated the data from 400 trials into a training set (300 trials) and a test set (100 trials). All trials received a near-threshold (50% detection rate) stimulus input to population E1 of area V1. The combined training and test set contained 200 hit and 200 miss trials, and these were randomly shuffled and allocated to the training and test sets.

As activity in region 9/46d was used to read out the trial outcome, we trained the classifier on activity in all other areas. Trials were considered a ’Hit’ if the mean activity in area 9/46d in the last 500ms before the end of the trial was greater than 15Hz, and a ’Miss’ otherwise. We trained each support vector classifier using scikit-learn in Python and standard parameter settings (Pedregosa et al. 2011). A separate classifier was trained for each timepoint in the training data. We then used each of these classifiers to predict the trial outcome based on activity at each time point in a separate test set. Finally, we compared these predictions to the actual trial outcome (defined according to the late sustained activity in 9/46d).

To estimate whether the coding pattern is similar between times *t* and *t*^*′*^, we can train a classifier at time *t* (across trials) and test it at time *t*^*′*^. When applied across all pairs of timepoints, this leads to a square *T* × *T* temporal generalization matrix, where *T* is the number of timepoints (King and Dehaene 2014; King et al. 2016; Meyers et al. 2008).

We assessed the strength of correlation between decoder coefficients (for the decoder trained at each timepoint) and the cortical hierarchy using Pearson correlations (Fig 3E). We conservatively judged the correlation at a particular timepoint to be significant only if the p value was less than 0.001 for all timepoints within a 10ms period centered on the timepoint.

### In-vitro receptor autoradiography

Quantitative in-vitro receptor autoradiography was applied to determine the densities of NMDA and receptors in cytoarchitectonically identified cortical areas of the macaque monkey brain (Impieri et al. 2019; Niu et al. 2020, 2021; Rapan et al. 2021, 2022a,b).

Brain tissue was shock frozen at -40°C in isopentane, hemispheres serially sectioned in the coronal plane at 20µm by means of a cryomicrotome, and sections thaw mounted onto glass slides. Alternating sections were processed for the visualization of cell bodies (Merker 1983) or of receptor densities according to previously published established protocols (Palomero-Gallagher and Zilles 2018; Table 1). In short, receptor incubation protocols consisted of a preincubation to rehydrate sections and remove endogenous ligands, a main incubation, and a washing step to stop the binding process and remove surplus ligand and buffer salts. The main incubation encompassed parallel experiments to identify the total and non-specific binding of each ligand, whereby sections were incubated with the radiolabeled ligand alone or with the radiolabeled ligand in conjunction with a non-labelled displacer, respectively.

**Table 1:** Incubation protocols

| Table 1 | AMPA | NMDA |
| --- | --- | --- |
| [<sup>3</sup>H]-Ligand | AMPA (10 nM) | MK-801 (3.3 nM) |
| Displacer | quisqualate (10 μM) | MK-801 (100 μM) |
| Incubation buffer | 50mM Tris-acetate (pH 7.2)<br>+ 100 mM KSCN* | 50mM Tris-acetate (pH 7.2)<br>+ 50 μM glutamate<br>+ 30 μM glycine*<br>+ 50 μM spermidine* |
| Preincubation | 3 x 10 min, 4°C | 15 min, 4°C |
| Main incubation | 45 min, 4°C | 60 min, 22°C |
| Final rinsing | 1. 4 x 4 sec, 4°C<br>2. 2 x 2sec in 100/2.5<br>acetone/glutaraldehyde, 4°C | 1. 2 x 5 min, 4°C<br>2. rinse in distilled water,<br>22 °C |

\* substance only included in buffer for the main incubation

**Table 2.** Parameters for Numerical Simulations

| Parameter | Description | Value |
| --- | --- | --- |
| $\tau^{NMDA}, \tau^{AMPA}$ | E synaptic time constants | 60ms, 2ms |
| $g^{NMDA}, g^{AMPA}$ | Channel Conductances | 1000pA, 10000pA |
| $\tau^{GABA}$ | I synaptic time constant | 5ms |
| $\tau^R$ | Firing rate time constant | 2ms |
| $\gamma^{NMDA}, \gamma^{AMPA}, \gamma^I$ | Synaptic rise constants | 1.282, 2, 2 |
| $\kappa^{sup}, \kappa^{dp}, \kappa^{local}$ | NMDA fraction | 0., 0.8, 0.91 |
| $\rho^{sup}, \rho^{dp}$ | Long-range E/I targets | 1., 0.015 |
| $z^{min}$ | Min excitation value | 0.6 |
| $z_I^{min}$ | Min excitation value | 0.218 |
| $\sigma_{noise}$ | std. dev. of noise | 2.5pA |
| $I_E^{bg}, I_I^{bg}$ | Background inputs | 329.4pA, 260pA |
| $a, b, d$ | f-I curve (E cells) | 0.135 Hz/pA, 54Hz, 0.308s |
| $\beta_i, I_{th}$ | f-I curve (I cells) | 153, 75Hz/nA, 252Hz |
| $b_1$ | Rescale FLN | 0.3 |
| $G_{E,E}^{N,loc}$ | Excitatory NMDA strengths | 480pA |
| $G_{E,E}^{A,loc}$ | Excitatory AMPA strengths | 4800pA |
| $G_{I,E}^{N,loc}$ | Excitatory NMDA strengths to the Inhib Unit | 10pA |
| $g_{E,I}, g_{I,I}$ | Inhibitory strengths | -8800pA, -120pA |
| $G_E^{NMDA}, G_I^{NMDA}$ | Long-range NMDA strength | 1500pA, 10.5pA |
| $G_E^{AMPA}, G_I^{AMPA}$ | Long-range AMPA strength | 15000pA, 105pA |
| $G_0$ | Local balanced coupling | 215pA |
| $I^{stim}$ | Stimulus strength | 250pA |

Please note that this is a current-based model, so all synaptic strengths area given in units of pA.

Radiolabelled sections were co-exposed with plastic standards calibrated to account for total brain protein content and with known concentrations of radioactivity against tritium (3H) sensitive films. The ensuing autoradiographs were digitized with an 8-bit grey value resolution for densitometric analysis (Palomero-Gallagher and Zilles, 2018). Hereby, calibration curves computed by non-linear, least-squares fitting were used to define the relationship between gray values and concentrations of radioactivity derived from the plastic standards. Radioactivity concentrations (*R*; in counts per minute, cpm) were converted to binding site concentrations (*Cb*; in fmol/mg protein) using the following equation:

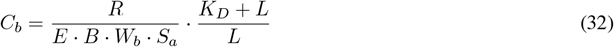

Where *E* is the efficiency of the scintillation counter, *B* is a constant representing the number of decays per unit of time and radioactivity (Ci/min), *Wb* the protein weight of a standard (mg), *Sa* the specific activity of the ligand used to label the target receptor (Ci/mmol), *K*_*D*_ the dissociation constant of the ligand (nM), and *L* the concentration of the ligand in the main incubation buffer (nM; determined by scintillation). Thus, the gray value of each pixel in a receptor autoradiograph could be transformed into a receptor density in fmol/mg protein. The location and extent of each cytoarchitectonically identified area was transferred to the neighboring autoradiographs and, for each receptor type separately, the mean (averaged over all cortical layers) of the grey values contained in 3-5 sections of the area in question was transformed into a receptor concentration per unit protein (fmol/mg protein).

### Receptor data-based model

For the receptor data-based model, we matched the total NMDA fraction to that seen in the data, adjusting for a constant mean shift between the model and receptor data, which we assume to be due to unmodelled connections (e.g. background inputs).

We calculate the overall NMDA fraction *X*_[*k*],*model*_(fraction of NMDA receptors over total number of excitatory receptors) in each area of the model.

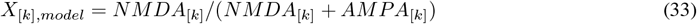

where *NMDA*_*k*_ and *AMPA*_*k*_ are the total local and inter-areal connections mediated by each receptor type. Here

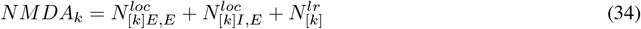

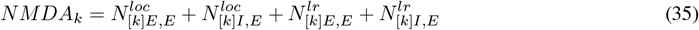

With:

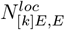 the number of NMDA receptors on the excitatory neurons coming from local connections.

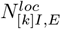 the number of NMDA receptors on the inhibitory neurons coming from local connections.

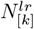 the total number of NMDA receptors coming from long-range connections.

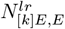 the number of NMDA receptors on the excitatory neurons coming from long-range connections.

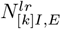 the number of NMDA receptors on the inhibitory neurons coming from long-range connections.

In the model, 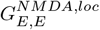 and 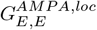 are set as follow:

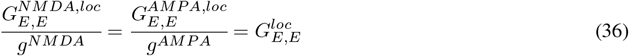

With *g*^*NMDA*^ the conductance due to one bound NMDA receptor and *g*^*AMP A*^ the conductance due to one bound AMPA receptor. We also define:

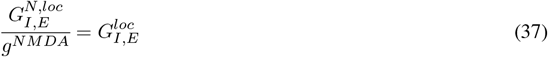

This equations can be expanded based on Equations (7), (10), (14) and (17)

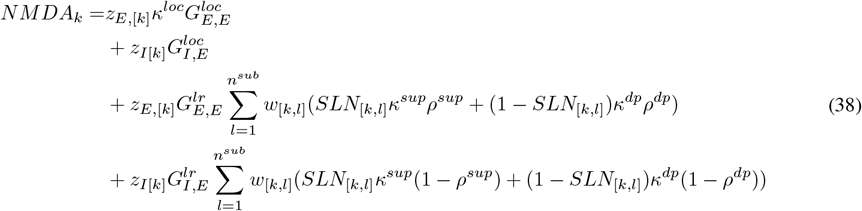

In the reference model, used throughout Figures 1-7, *κ*^*loc*^ is the same across all cortical areas.

Similarly:

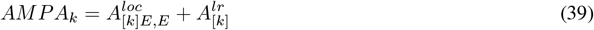

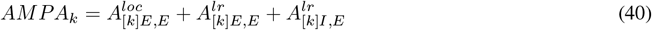

It is worth noting that their are no local AMPA connections targeting the inhibitory pool.

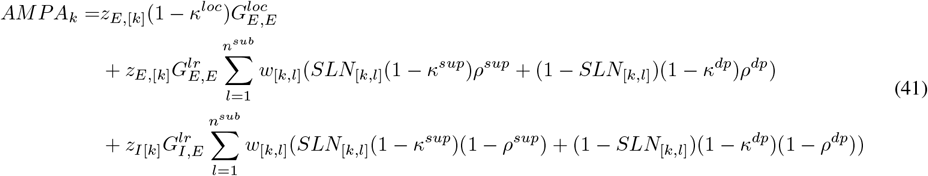

In practice, both *NMDA*_*k*_ and *AMPA*_*k*_ should be doubled (to represent the two excitatory populations), but as this affects all terms in the numerator and denominator, it will not affect the fraction *X*_[*k*]_.

We show in Figure 2 how the proportion of superficial and deep layer projections mediated by AMPA and NMDA receptors should lie in a particular range in order to enable rapid ‘ignition’ of cortical activity. Therefore, for the receptor data-based model, we treat these long-range feedforward and feedback NMDA and AMPA fractions as fixed. We then set the overall NMDA fraction in each area to match the experimentally-measured value *X*_[*k*],*data*_, shifted by a constant term *c* to account for the a mean shift between the raw receptor data and the reference model used in the rest of the paper. We can then calculate the local NMDA fraction 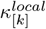 in each area required to match the observed NMDA fraction distribution across the cortex.

By reorganising equations 33 - 41, we can compute the local fraction 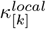 as a function of network parameters and real receptor data *X*_[*k*],*data*_. We forced 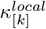 to lie between 0 and 1 using a clip function.

## Notes

### Competing Interest Statement

The authors have declared no competing interest.

### Summary of Updates

The manuscript has been re-worked to be easier to read and focus more on the detection dynamics.

