## Supplementary Figures for "A mesoscale connectome-based model of conscious access in the macaque monkey"

October 29, 2024

### Supplementary Figures

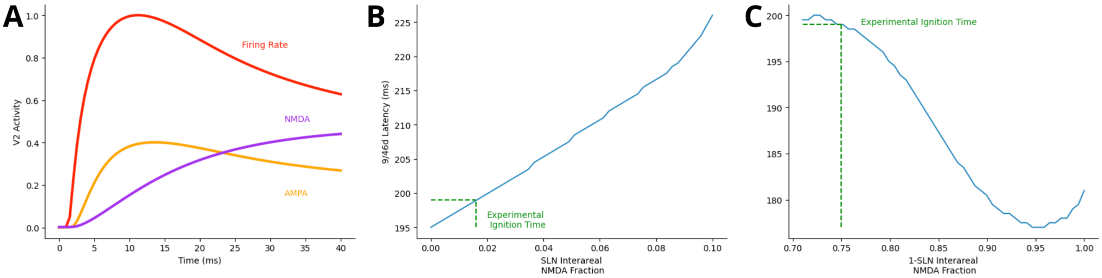


**Supplementary Figure 1.** *A) The dynamics of activity in area V2 (normalized firing rate) reflects the distinct dynamics of AMPA and NMDA receptors at feedforward and feedback connections. B) Impact of Receptor Type on Ignition Timing. Ignition times (time to reach 95% of peak dlPFC activity) vary with the fraction of NMDA receptors in inter-areal connections from B) SLN projections and C) 1-SLN projections.*

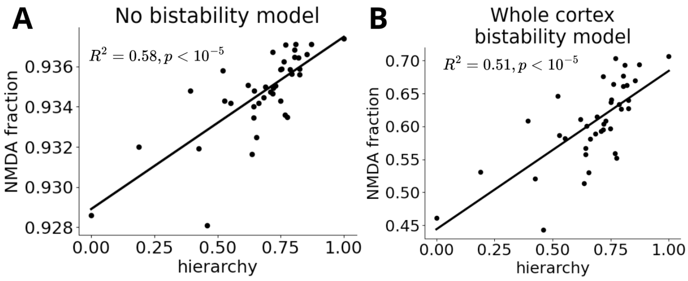


**Supplementary Figure 2.** *For both alternative models, the fraction of excitatory inputs via NMDA receptors (compared to total NMDA+AMPA inputs) increases along the cortical hierarchy.*
